# Intranuclear polyglycine aggregation drives neurodegeneration through epigenetic repression of chromatin accessibility and transcription

**DOI:** 10.64898/2026.05.11.723935

**Authors:** Yangye Lian, Shaoping Zhong, Jiaxin Huang, Luyao Huang, Yuzhe Li, Jingzhen Liang, Xin Wang, Jing Ding

## Abstract

Polyglycine (polyG) proteins translated from expanded GGC trinucleotide repeats are implicated in a growing group of neuromuscular degenerative disorders characterized by intranuclear inclusions, yet the pathogenic importance of aggregate localization and the mechanisms underlying polyG-induced neurodegeneration remain unclear. Here we show that intranuclear polyG aggregates are markedly more pathogenic than cytoplasmic aggregates in cellular and mouse models. Intranuclear aggregation causes greater cell death, more severe behavioral deficits and neuropathology, and earlier mortality. Mechanistically, intranuclear polyG aggregates impair nascent RNA synthesis and are associated with a transcriptionally repressive chromatin state marked by globally reduced chromatin accessibility, decreased H3K27 acetylation, and increased HDAC3 expression across cellular, mouse, and human disease tissue. Using a light-inducible system, we further show that this transcriptional impairment depends on insoluble intranuclear aggregate formation rather than diffuse polyG alone. Pharmacological HDAC inhibition partially restores histone acetylation and transcriptional output and ameliorates behavioral and pathological abnormalities. Together, these findings identify intranuclear polyG aggregates as the more pathologically relevant species and uncover epigenetic repression of chromatin accessibility and transcription as a potentially common mechanism underlying polyG diseases.

## INTRODUCTION

Nucleotide repeat expansions are associated with more than 50 human neurological disorders(1). Multiple pathogenic mechanisms have been proposed, encompassing transcriptional, translational, and post-translational dysregulation(2). In particular, transcribed repeats in coding or non-coding regions can give rise to repeat-containing proteins that exert toxic gain-of-function effects. Prototypical examples include the production of mutant huntingtin with expanded polyglutamine (polyQ) tracts in Huntington disease and dipeptide repeat proteins (DRPs) in *C9ORF72*-associated amyotrophic lateral sclerosis and frontotemporal dementia (ALS/FTD)(3). These aberrant protein products are highly aggregation-prone, readily forming inclusion bodies and disrupting a wide range of cellular processes, although their downstream pathogenic consequences remain incompletely understood in most conditions.

Recently, GGC trinucleotide repeat expansions at various genetic loci have been revealed to cause a group of neurodegenerative and neuromuscular diseases with clinical and radiological overlaps(4). These include fragile X-associated tremor/ataxia syndrome (FXTAS), neuronal intranuclear inclusion disease (NIID), oculopharyngeal myopathy with leukoencephalopathy (OPML), and oculopharyngodistal myopathies (OPDMs), caused by GGC expansions (60-250 repeats typically) in *FMR1*, *NOTCH2NLC*, *NUTM2B-AS1*, and *LRP12/GIPC1/NOTCH2NLC/RILPL1*, respectively. A shared pathological hallmark is the presence of eosinophilic p62- and ubiquitin-positive intranuclear inclusions in multiple cell types(5, 6). These similarities strongly suggest a common underlying pathogenic mechanism.

Accumulating evidence, including findings from our own group, has demonstrated that despite residing in the untranslated regions, GGC expansions are translated into polyglycine (polyG)-containing short peptides that are aggregation-prone and flanked by locus-specific amino acid residues, i.e. FMRpolyG, uN2CpolyG, uGIPC1polyG, asRILpolyG, and LOC6polyG(7–13). These observations led to the proposal of a unified disease spectrum termed polyG diseases(11, 14, 15). Prior studies further indicate that polyG-containing peptides determine the inclusion formation and mediate cellular toxicity through diverse mechanisms, such as disruption of the nuclear lamina(10), altered nucleocytoplasmic transport(9, 11), disturbed mitochondrial function(16), and dysregulated mRNA(12) or tRNA splicing(17). Although some of these pathogenic effects have been attributed to the flanking sequences adjacent to the polyG tracts in *FMR1* or *NOTCH2NLC*(^9, 10^), emerging evidence increasingly supports the expanded polyG tracts themselves as the principal toxic species across diseases(17, 18).

A distinct feature of polyG diseases lies in their striking predominance of intranuclear inclusions in human tissues. Only a few studies have reported rare and suspicious cytoplasmic inclusions with atypical morphologies in human fibroblasts or muscles from NIID patients(19, 20). This feature contrasts with polyQ diseases and *C9ORF72*-associated ALS/FTD, in which both nuclear and cytoplasmic inclusions are observed routinely(21, 22). However, in current animal models overexpressing polyG-containing peptides, cytoplasmic inclusions are abundant and often more prevalent than intranuclear inclusions(9–12, 18). This discrepancy raises a critical question as to whether cytoplasmic inclusions observed in experimental models faithfully reflect the pathophysiological mechanisms operating in humans. Furthermore, the pathological consequences and mechanistic significance of polyG in different subcellular compartments remain poorly understood.

In this study, we sought to determine whether the subcellular localization of polyG aggregates influences their pathogenicity and to define the molecular mechanisms underlying polyG-induced neurodegeneration in mouse models and human tissues. Our study identified that intranuclear polyG aggregates are markedly more toxic than cytoplasmic aggregates and suppress nascent RNA synthesis, accompanied with reduced chromatin accessibility, decreased H3K27 acetylation (H3K27ac), and increased HDAC3 expression. These findings suggest that intranuclear aggregation is critical for polyG-induced neurodegeneration and uncover epigenetic repression of chromatin accessibility and transcription as a common mechanism of polyG diseases.

## RESULTS

### Construction of polyG fusion proteins with controllable subcellular localization

To examine the toxicity of the polyG tract itself, independent of flanking sequences or repeat RNA toxicity, we developed a construct comprising 73 glycine residues encoded by GGN repeats (N represents A/G/T/C) and followed by a FLAG tag (Supplemental Figure 1A). Overexpressing this construct in SH-SY5Y cells induced the formation of both intranuclear and extranuclear aggregates. Adding nuclear localization sequence (NLS) to this construct did not alter the subcellular distribution pattern significantly (Supplemental Figure 1, B and C), which may be attributed to the ability of small proteins to passively diffuse through nuclear pore complexes (NPCs).

As prior studies demonstrated that EGFP homomultimers hinder their passive diffusion through NPC with increasing molecular weight(23, 24), we constructed a series of constructs fused with 1× or 2×EGFP and NLS, confirmed by Sanger sequencing (Supplemental Figure 1A, 2A). The proportion of intranuclear aggregates was significantly increased (57.2%) in the 1×EGFP-NLS construct, and even higher (80.8%) in the 2×EGFP-NLS construct (Supplemental Figure 1, B and C). Replacing NLS with NES induced a remarkably reduced intranuclear localization (6.6%). These findings indicate that increasing the size of the fusion protein markedly improved experimental control over inclusion localization.

The 2×EGFP-based constructs were selected for subsequent experiments and designated as Gly73, Gly73-NLS, and Gly73-NES, together with the short polyG control Gly7 (Figure 1A). Representative fluorescence imaging confirmed that Gly73-NLS predominantly formed intranuclear aggregates, whereas Gly73-NES mainly formed extranuclear aggregates, with Gly73 showing an intermediate distribution (Figure 1, B and C). These aggregates co-localized with p62, consistent with the pathological findings observed in patients with polyG-related disorders (Supplemental Figure 2E). We next examined their mRNA and protein abundance. The mRNA level of Gly7 was higher than that of the polyG-containing constructs in the qPCR assay, whereas Gly73, Gly73-NLS, and Gly73-NES were expressed at broadly comparable transcript levels (Supplemental Figure 2B). Immunoblotting further showed that Gly73-NLS exhibited somewhat lower protein abundance than Gly73 and Gly73-NES (Supplemental Figure 2, C and D).

**Figure 1.**
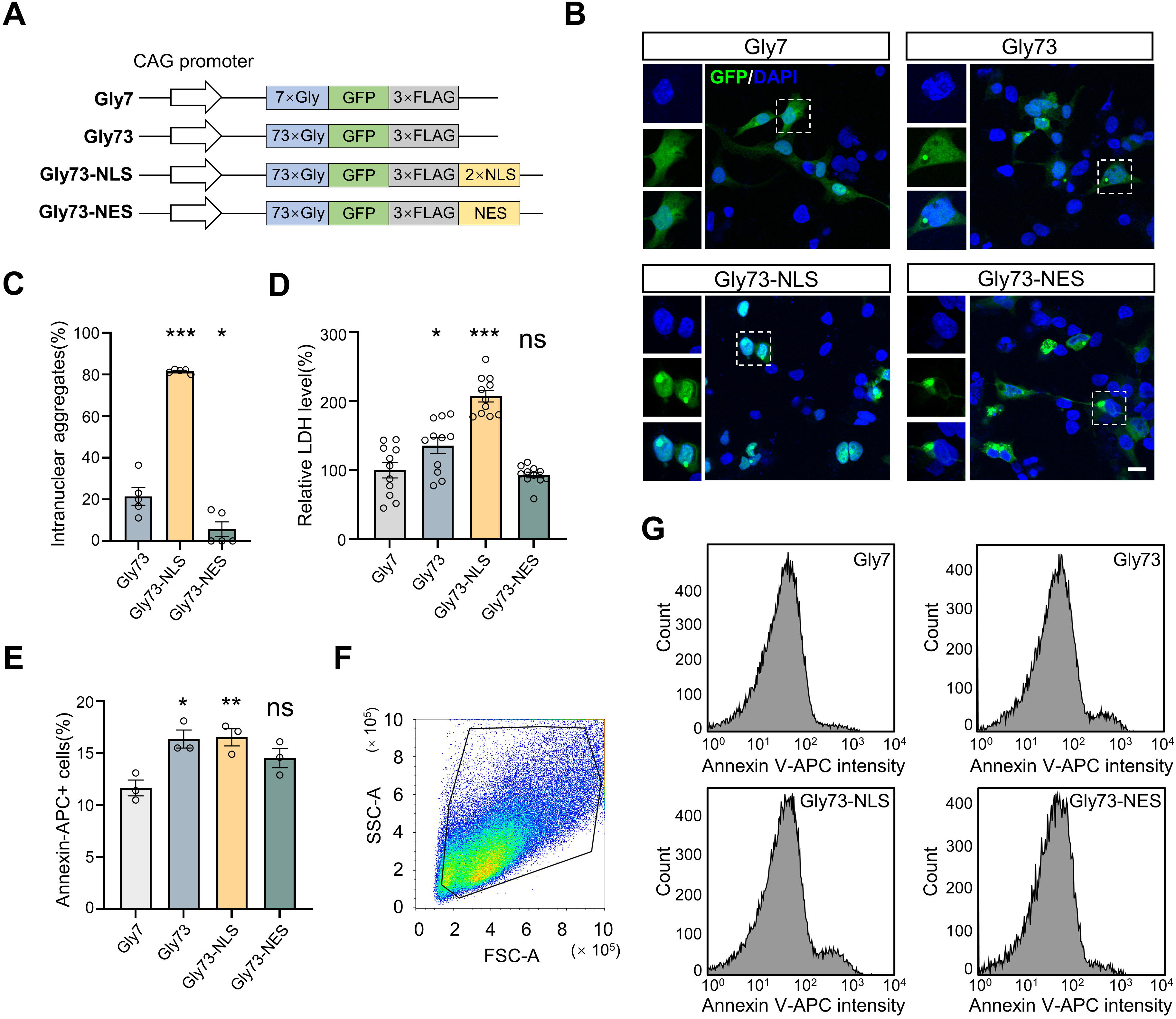
Intranuclear polyG aggregation causes enhanced cytotoxicity and cell death. **(A)** Schematic of the polyG constructs used in this study. Gly7 served as a short-repeat control, whereas Gly73 contained an expanded 73-glycine tract. To bias the subcellular distribution of aggregates, Gly73 was fused to either a 2×NLS or an NES, generating Gly73-NLS and Gly73-NES, respectively. All constructs were expressed under the CAG promoter and fused to 2×EGFP and a 3×FLAG tag. **(B)** Representative confocal images of SH-SY5Y cells expressing the indicated constructs. Gly73-NLS predominantly formed intranuclear aggregates, whereas Gly73-NES mainly formed extranuclear aggregates, with Gly73 showing an intermediate distribution. Boxed regions are shown at higher magnification. Scale bar, 50 μm. **(C)** Quantification of the percentage of intranuclear aggregates among total aggregates in cells expressing the indicated constructs (n=5). **(D)** LDH release assay showing cytotoxicity induced by the indicated constructs in SH-SY5Y cells (n=11). **(E)** Quantification of Annexin V-APC-positive cells by flow cytometry, showing increased apoptosis in the Gly73-NLS group (n=3). **(F)** Representative flow cytometry gating plots used for Annexin V-APC analysis. **(G)** Representative histograms of Annexin V-APC fluorescence intensity in cells expressing the indicated constructs. Data are means ± SEM and analyzed with one-way ANOVA (**C-E**). ‘‘ns’’ represents non-significant; *p < 0.05, **p < 0.01, and ***p < 0.001.

### Intranuclear polyG aggregation induces cytotoxicity and cell death

We overexpressed the indicated constructs in SH-SY5Y cells and measured lactate dehydrogenase (LDH) release into the culture medium as an indicator of cell death and injury. LDH levels were significantly increased in the Gly73-NLS group compared with the Gly7 group, whereas the Gly73-NES group showed no significant increase (Figure 1D). Apoptosis was assessed by flow cytometric analysis of Annexin V-APC staining. Consistent with the LDH results, the proportion of Annexin V-positive cells was significantly higher in the Gly73-NLS group than in the Gly7 control group, whereas the Gly73-NES group did not differ significantly from the Gly73 group (Figure 1, E-G). Together, these findings indicate that intranuclear polyG aggregates are associated with greater cytotoxicity and pro-apoptotic effects.

### Intranuclear polyG aggregation causes more severe neurodegenerative phenotypes

These constructs were packaged into AAVs of the PHP.eB serotype, known for efficient transduction throughout the mouse brain(25), and injected intravenously into wild-type C57BL/6 mice, followed by behavioral and survival analyses (Figure 2A). At 2 months after AAV injection, Gly73-NLS mice showed a significant reduction in body weight (Figure 2B). Motor tests, including the rotarod test, the notched bar test, the hindlimb extension test revealed motor impairment in all polyG-expressing groups, with the most severe deficits consistently observed in the Gly73-NLS group (Figure 2, C-F). By contrast, Gly7 control mice showed no overt motor abnormalities. We next examined cognitive function at 2.5 months after AAV injection. In the Y-maze test, spontaneous alternation was significantly reduced in the Gly73, Gly73-NLS, and Gly73-NES groups, with the largest decrease observed in the Gly73-NLS group (Figure 2G). In the novel object recognition test, Gly73-NLS mice traveled a significantly greater distance, suggesting increased irritability or hyperactivity-like behavior, and spent the least time exploring the novel object (Figure 2, H-J).

**Figure 2.**
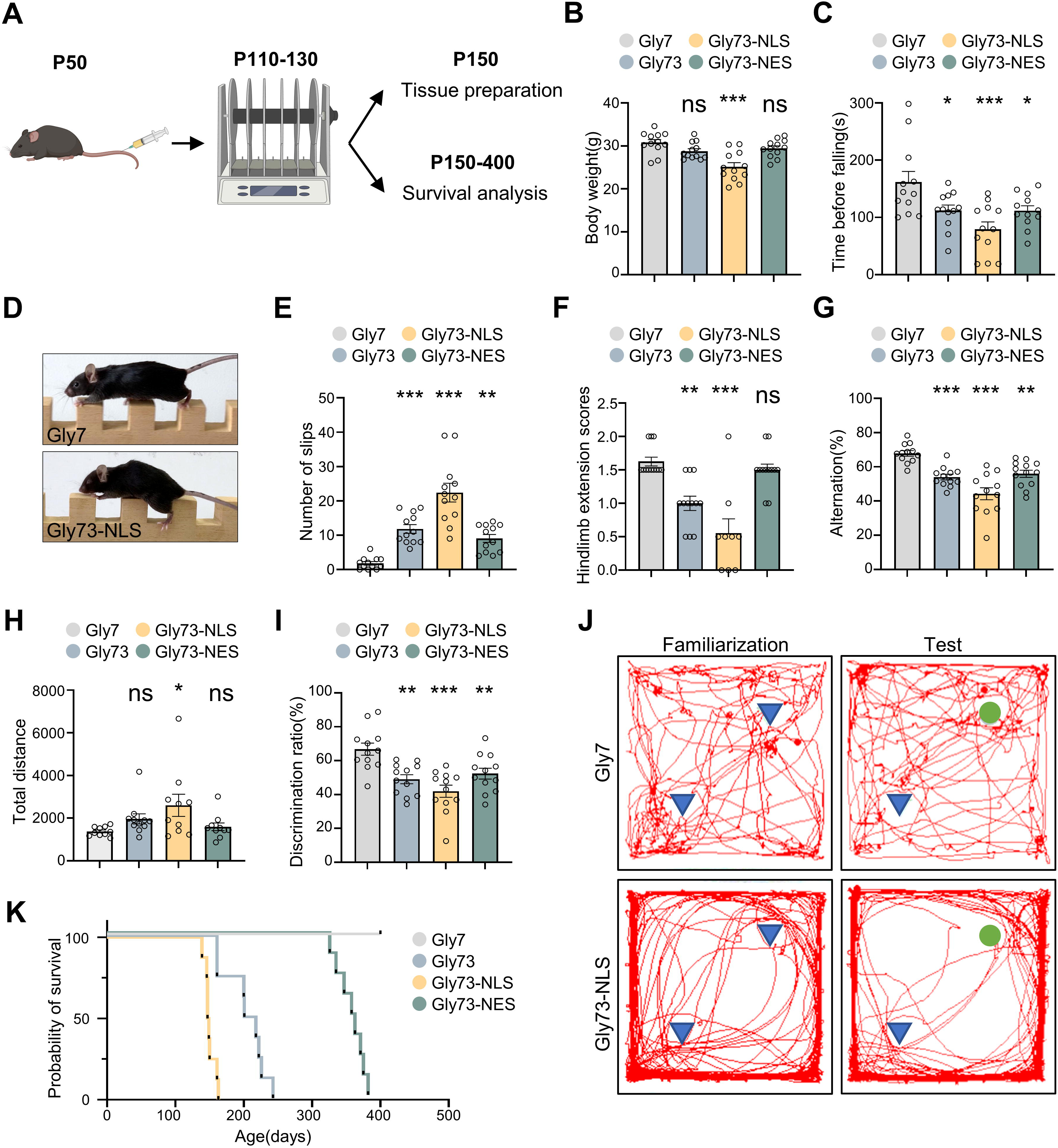
Intranuclear polyG aggregation causes the most severe survival, motor, and cognitive deficits in vivo. **(A)** Schematic of the experimental design. Mice received AAV injection at P50. Behavioral assessments were performed between P110 and P130, followed by tissue collection at P150 or long-term survival monitoring up to P150–400. **(B)** Body weight of mice at 2 months after AAV injection (n=12). **(C)** Rotarod performance shown as latency to fall (n=12). **(D)** Representative images from the notched bar test. **(E)** Quantification of slips in the notched bar test (n=12). **(F)** Hindlimb extension scores (n=9–12). **(G)** Y-maze spontaneous alternation (n=12). **(H)** Total travel distance in the novel object recognition test (n=10). **(I)** Discrimination ratio in the novel object recognition test (n=12). **(J)** Representative movement traces during the novel object recognition test in the familiarization and test phases. **(K)** Kaplan–Meier survival curves of mice. Data are means ± SEM and analyzed with one-way ANOVA (**B, C, E, F, G, H, I**) and log-rank (Mantel–Cox) test (**K**). ‘‘ns’’ represents non-significant; *p < 0.05, **p < 0.01, and ***p < 0.001.

Survival analysis showed marked differences among the groups. Mice in the Gly73-NLS group exhibited the earliest mortality, with deaths beginning approximately 3-4 months after injection. The Gly73 and Gly73-NES groups exhibited intermediate survival, with obvious deaths observed at 5-6 and 9-11 months after injection, respectively. No deaths were observed in the control group throughout the 14-month follow-up period (Figure 2K). Overall, these results indicate that mice in the Gly73-NLS group consistently displayed the most severe abnormalities across survival, motor, and cognitive assessments, followed by the Gly73 group, whereas the Gly73-NES group generally showed a milder phenotype.

### Intranuclear polyG aggregation induces more severe pathology in vivo

To determine the pathological consequences in vivo, we examined mouse brains at P150 after AAV injection. Intranuclear aggregates accounted for 77.2% of total polyG aggregates in the cerebral cortex of the Gly73-NLS group, compared with 32.6% in the Gly73 group and 10.7% in the Gly73-NES group (Figure 3, A and B). Transmission electron microscopy revealed abundant intranuclear aggregates in the cortex of Gly73-NLS mice, whereas such aggregates were rarely observed in Gly73-NES mice, in which aggregates were predominantly cytoplasmic. Ultrastructurally, intranuclear aggregates consisted of irregularly arranged filaments and occasionally contained a central electron-dense core, while cytoplasmic aggregates exhibited a dense central core surrounded by radially oriented filaments (Figure 3C). These ultrastructural features were consistent with our previous report about mice overexpressing uN2CpolyG(18).

**Figure 3.**
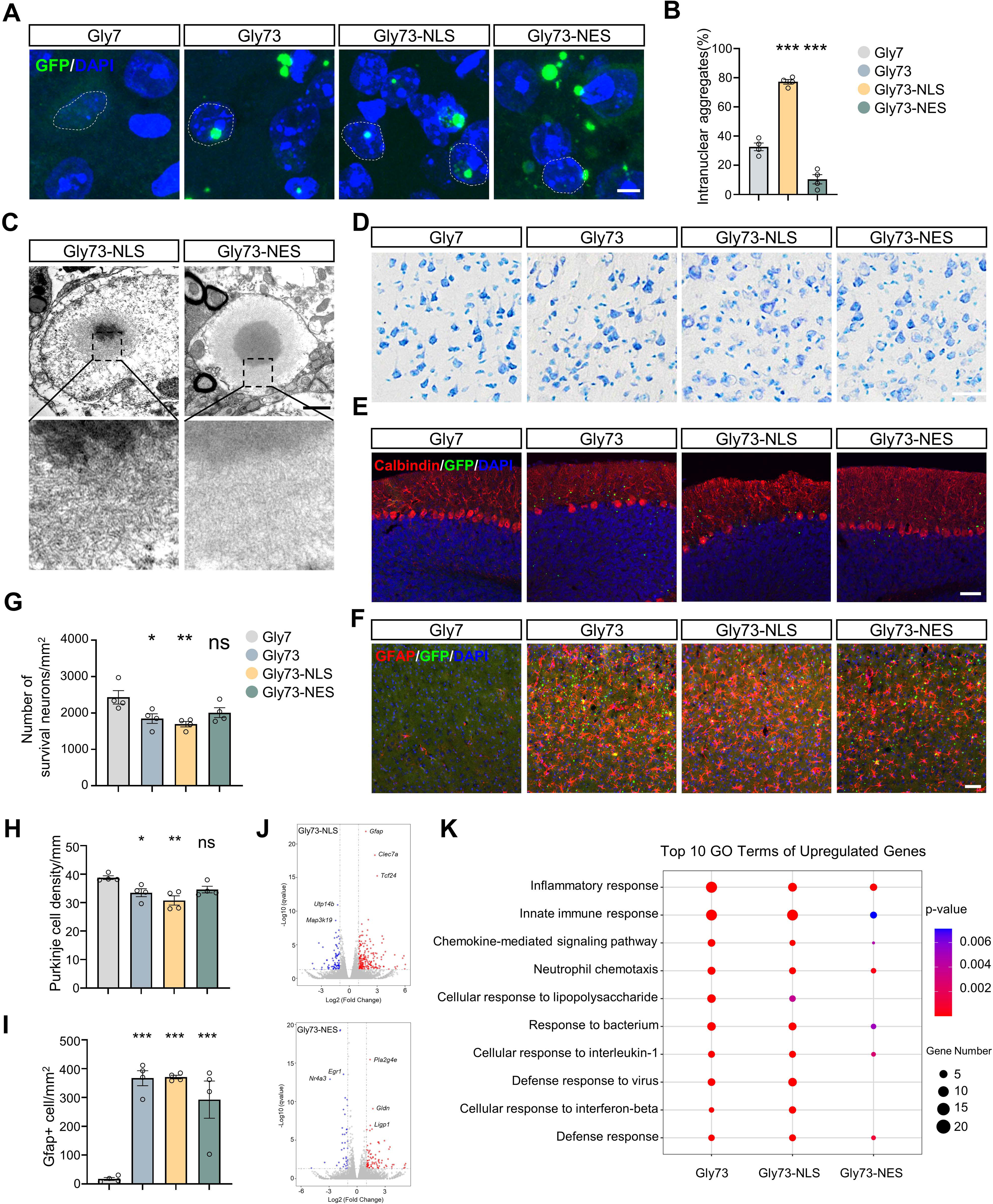
Intranuclear polyG aggregation induces severe neuropathology and inflammatory transcriptional responses in mouse brain. **(A)**Representative confocal images showing the subcellular distribution of GFP-positive aggregates in the cerebral cortex of mice expressing Gly7, Gly73, Gly73-NLS, or Gly73-NES. Dashed circles indicate nuclei. Scale bar, 5 μm. **(B)** Quantification of the percentage of intranuclear aggregates among total aggregates in the indicated groups (n=4). **(C)** Transmission electron microscopy images showing representative intranuclear aggregates in Gly73-NLS mice and cytoplasmic aggregates in Gly73-NES mice. Insets show higher-magnification views of the boxed regions. Scale bars, 1 μm. **(D)** Representative Nissl staining images of the cerebral cortex from mice of the indicated groups at P150. Scale bar, 50 μm. **(E)** Representative calbindin staining images showing Purkinje cells in the cerebellum of the indicated groups at P150. Scale bar, 50 μm. **(F)** Representative GFAP immunostaining images in the cerebral cortex of the indicated groups at P150. Scale bar, 50 μm. **(G)** Quantification of surviving cortical neurons based on Nissl staining (n=4). **(H)** Quantification of calbindin-positive Purkinje cell density (n=4). **(I)** Quantification of GFAP-positive cell density (n=4). **(J)** Volcano plot of gene expression changes in the Gly73-NLS and Gly73-NES groups compared to the Gly7 group. **(K)** Gene Ontology analysis of upregulated genes in the indicated groups, showing enrichment of inflammatory and innate immune pathways. Data are means ± SEM and analyzed with one-way ANOVA (**B, G, H, I**). ‘‘ns’’ represents non-significant; *p < 0.05, **p < 0.01, and ***p < 0.001.

Beyond inclusion pathologies, Nissl staining showed a reduction in surviving cortical neurons in Gly73-NLS mice (Figure 3, D and G), consistent with the observations in the NeuN immunostaining (Supplemental Figure 3, A and C). In the cerebellum, calbindin staining revealed significant Purkinje cell loss in Gly73 and Gly73-NLS mice, whereas the reduction in Gly73-NES mice was relatively mild and did not reach significance (Figure 3, E and H). GFAP and Iba1 immunostaining revealed prominent astrogliosis and microgliosis in Gly73 and Gly73-NLS mice, whereas the increase in Gly73-NES mice was comparatively modest (Figure 3, F and I; Supplemental Figure 3, B and D). Together, these results indicate that intranuclear polyG aggregates are associated with more severe neuronal pathology and robust glial activation.

### Molecular changes associated with intranuclear polyG aggregation

To define the molecular consequences of nuclear versus cytoplasmic polyG aggregation, we performed bulk RNA-seq on the cerebral cortex at P150. PolyG expression induced broad transcriptional changes relative to Gly7 controls, with the largest number of differentially expressed genes detected in the Gly73-NLS group (Figure 3J; Supplemental Figure 3E). Consistent with the prominent gliosis, upregulated genes in Gly73-NLS cortices were strongly enriched for inflammation- and innate immunity-related pathways, including chemokine signaling, cytokine-responsive programs, and antiviral defense. Representative upregulated genes included *Gfap*, *Clec7a*, *ligp1* and *Trem2*. Similar but weaker enrichment was observed in the Gly73-NES group (Figure 3K). These data indicate that nuclear polyG aggregation is associated with broader transcriptional perturbation and a stronger neuroinflammatory response than cytoplasmic aggregation.

### Intranuclear polyG aggregation disrupts nascent RNA synthesis

Because intranuclear polyG aggregates exhibited greater toxicity than cytoplasmic aggregates, we next examined whether essential nuclear functions were impaired. Given the central importance of transcriptional homeostasis for neuronal survival and function, we assessed nascent RNA synthesis by 5-ethynyl uridine (EU) labeling in cells expressing the indicated constructs. Compared with Gly7 controls, Gly73-NLS expression caused a marked reduction in EU incorporation, whereas the decreases in the Gly73 and Gly73-NES groups were less pronounced, indicating that nuclear targeting of polyG was associated with stronger suppression of nascent RNA synthesis (Figure 4, A and B). To determine whether a similar defect occurs in vivo, we administered EU to mice at P150 and assessed EU incorporation in the cerebral cortex. Consistent with our in vitro findings, cortical neurons from Gly73-NLS mice showed a significant reduction in EU intensity compared with Gly7 mice, whereas the decrease in the Gly73 and Gly73-NES groups was relatively modest (Figure 4, C and D).

**Figure 4.**
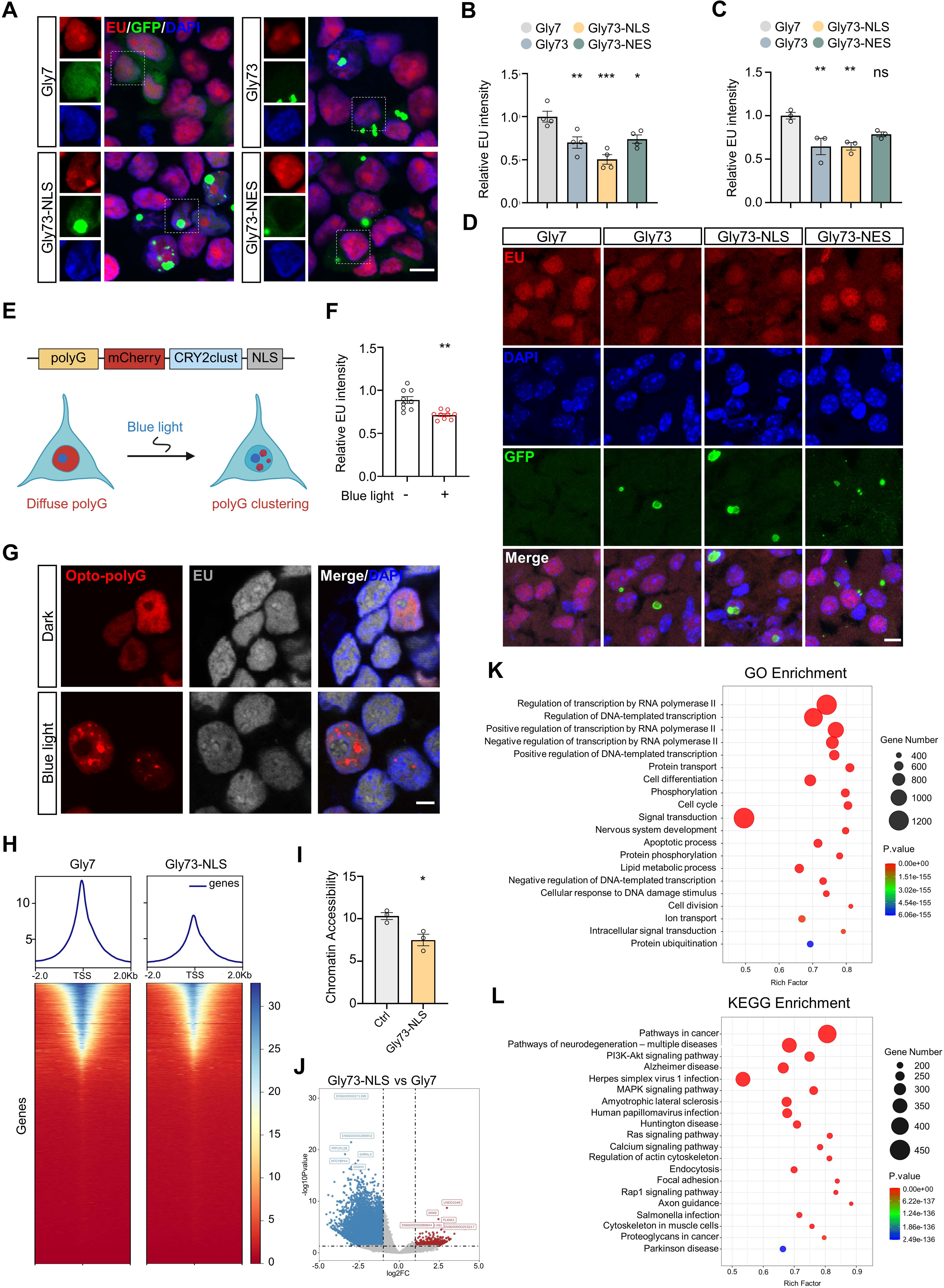
Intranuclear polyG aggregation impairs nascent RNA synthesis and is associated with reduced chromatin accessibility. **(A)** Representative images of 5-ethynyl uridine (EU) labeling in SH-SY5Y cells expressing Gly7, Gly73, Gly73-NLS, or Gly73-NES. Insets show enlarged views of the boxed regions. Scale bar, 10 μm. **(B)** Quantification of relative EU intensity in cells expressing the indicated constructs (n=4). **(C)** Quantification of relative EU intensity in the cerebral cortex of the indicated groups (n=3). **(D)** Representative images of EU labeling in the cerebral cortex of mice expressing Gly7, Gly73, Gly73-NLS, or Gly73-NES at P150. Scale bar, 10 μm. **(E)** Schematic diagram of the optogenetic system. The fusion protein consists of Cry2, mCherry, and polyG. **(F)** Quantification of relative EU intensity in the optogenetic polyG system under dark or blue-light conditions (n=9). **(G)** Representative images of the optogenetic polyG system under dark or blue-light conditions. Blue-light illumination induces formation of intranuclear opto-polyG aggregates. Scale bar, 5 μm. **(H)** TSS-centered accessibility profiles and heatmaps showing reduced open chromatin signals in Gly73-NLS cells relative to Gly7 controls. **(I)** Global chromatin accessibility measured by ATAC-seq in SH-SY5Y cells expressing Gly7 or Gly73-NLS (n=3). **(J)** Volcano plot showing differentially accessible regions in Gly73-NLS cells relative to Gly7 controls. **(K)** Gene Ontology enrichment analysis of genomic regions with reduced accessibility in Gly73-NLS cells. **(L)** KEGG pathway enrichment analysis of genomic regions with reduced accessibility in Gly73-NLS cells. Data are means ± SEM and analyzed with an unpaired two-tailed Student’s t test (**F, G**) and one-way ANOVA (**B, C**). ‘‘ns’’ represents non-significant; *p < 0.05, **p < 0.01, and ***p < 0.001.

We next asked whether this reduction in nascent RNA synthesis was accompanied by changes in the steady-state mRNA pool. Oligo(dT) staining revealed reduced mRNA signal in cells expressing Gly73-NLS, whereas the changes in the Gly73 and Gly73-NES groups were less evident (Supplemental Figure 4, A and B). Together, these findings indicate that intranuclear polyG aggregation suppresses nascent RNA synthesis and reduces global mRNA levels.

### Aggregation is essential for polyG-induced nascent RNA synthesis disruption

We next asked whether the global reduction in nascent RNA synthesis specifically depended on the aggregation of intranuclear polyG, rather than merely reflecting the presence of soluble polyG. To address this, an optogenetic polyG system was developed based on CRY2clust, a blue-light-responsive clustering module that enables rapid and reversible protein oligomerization(26, 27). We generated a Gly49-mCherry-CRY2clust-NLS (Opto-polyG) construct to inducibly form polyG aggregates in the nucleus (Figure 4E). Gly49 was selected because polyG aggregation increases with repeat length, whereas our previous work showed that Gly49 has relatively weak intrinsic aggregation propensity, thereby minimizing spontaneous aggregate formation before light stimulation and improving temporal control(11).

Using this system, we found that blue-light-induced intranuclear Opto-polyG aggregation significantly reduced EU incorporation compared with the dark condition (Figure 4, F and G). Consistent with this result, in cells expressing the Gly73-NLS construct, cells harboring visible aggregates exhibited significantly lower EU intensity than aggregate-negative cells showing only diffuse GFP signal (Supplemental Figure 4, C and D). Moreover, EU signal was inversely correlated with aggregate size (Supplemental Figure 4E), supporting a burden-dependent inhibitory effect of intranuclear aggregates on nascent RNA synthesis. Together, these results indicate that polyG-mediated suppression of RNA synthesis requires not only nuclear localization, but also aggregation.

### Intranuclear polyG aggregation induces globally reduced chromatin accessibility

The reduction in nascent RNA synthesis prompted us to investigate upstream regulatory mechanisms that might constrain transcriptional output. Because chromatin accessibility is a key determinant of active transcription(28), we performed ATAC-seq in SH-SY5Y cells expressing Gly73-NLS or Gly7 (Supplemental Figure 5A). ATAC-seq revealed a global reduction of chromatin accessibility in Gly73-NLS cells relative to Gly7 controls (Figure 4I). Transcription start site (TSS)-centered accessibility profiles and heatmap visualization showed broadly attenuated open chromatin signals in the Gly73-NLS group, indicating a widespread loss of transcription-permissive chromatin states (Figure 4H; Supplemental Figure 5 C). Unsupervised clustering further separated Gly73-NLS samples from Gly7 controls, supporting a distinct chromatin landscape associated with intranuclear polyG aggregation (Supplemental Figure 5B).

Differential accessibility analysis showed that regions with reduced accessibility were markedly more abundant than regions with increased accessibility in Gly73-NLS cells (Figure 4J). Analysis of genomic distribution further showed that these accessibility changes were broadly distributed across multiple chromosomes and genomic features, including promoter-proximal and intragenic regions, rather than being restricted to a specific genomic compartment (Supplemental Figure 5, D and E). Functional enrichment analysis of the less accessible regions revealed significant overrepresentation of pathways related to transcriptional regulation, including RNA polymerase II-dependent transcription and DNA-templated transcription, as well as pathways linked to neuronal system function and apoptotic processes (Figure 4, K and L). Representative genome browser tracks further illustrated decreased accessibility at selected loci, including genes such as *ANKRD1* and *TGFBI*, in the Gly73-NLS group (Supplemental Figure 5, F and G). Together, these findings indicate that intranuclear polyG aggregation is associated with globally reduced chromatin accessibility.

### Intranuclear polyG aggregation induces decreased histone acetylation

The marked reduction in chromatin accessibility suggested the presence of repressive epigenetic remodeling. Because H3K27ac marks transcriptionally active and open chromatin regions(29, 30), we next asked whether this modification was altered in association with polyG aggregation. In SH-SY5Y cells, immunofluorescence staining showed that H3K27ac intensity was reduced in the Gly73-NLS group compared with Gly7 controls, whereas the changes in the Gly73 and Gly73-NES groups were less pronounced (Figure 5, A and B). We then extended this analysis to the mouse model. Western blotting analysis of brain lysates showed reduced H3K27ac intensity in the Gly73-NLS group (Figure 5, C and D) and this decrease was further confirmed by immunostaining of brain sections (Figure 5, E and G).

**Figure 5.**
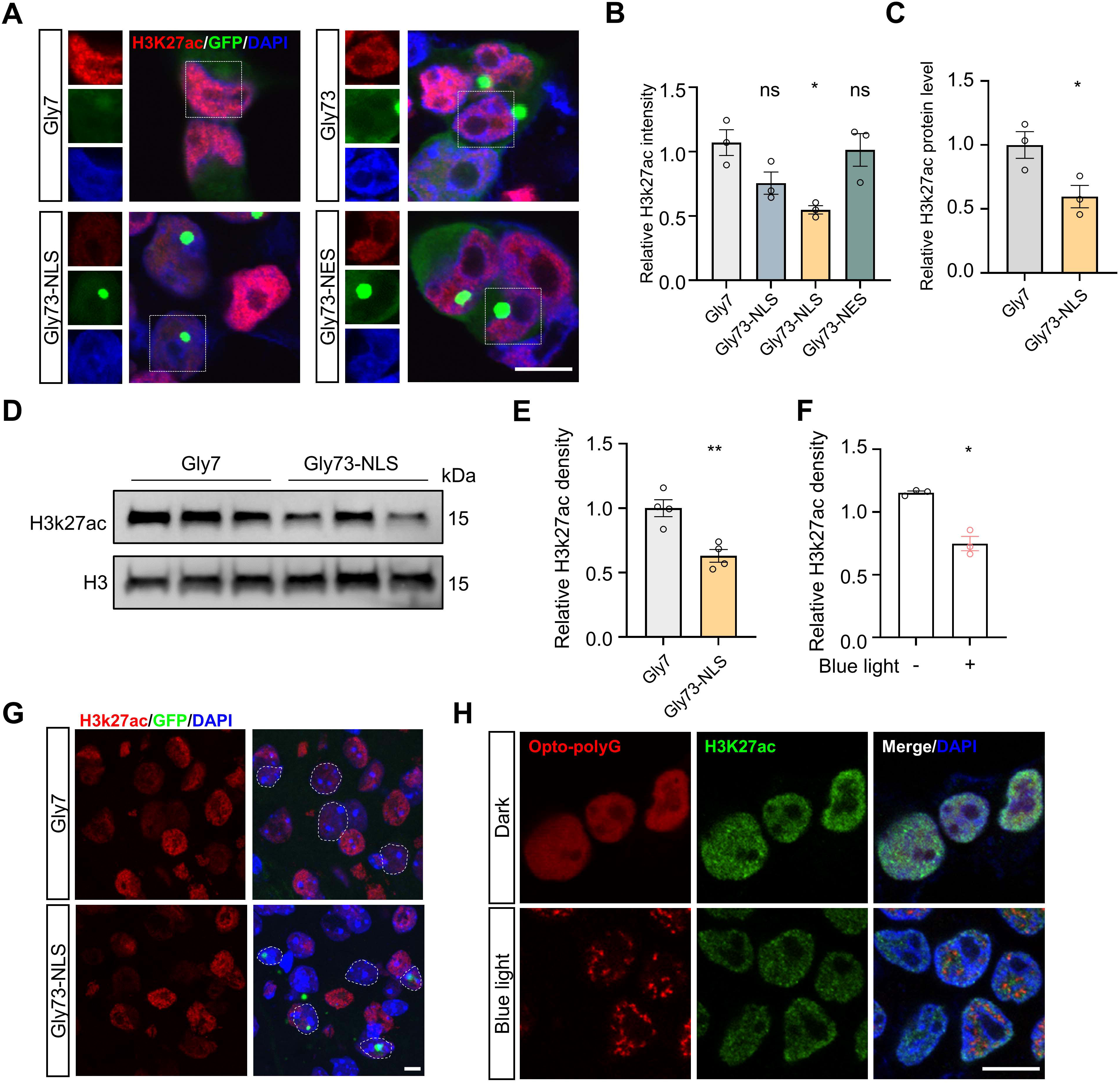
Intranuclear polyG aggregation induces decreased H3K27 acetylation in polyG models. **(A)** Representative images of H3K27ac immunostaining in SH-SY5Y cells expressing Gly7, Gly73, Gly73-NLS, or Gly73-NES. Scale bar, 10 μm. **(B)** Quantification of relative H3K27ac intensity in SH-SY5Y cells expressing the indicated constructs (n=3). **(C)** Quantification of relative H3K27ac protein levels shown in (**D**) (n=3). **(D)** Representative immunoblot of H3K27ac in brain lysates from Gly7- and Gly73-NLS-expressing mice. H3 served as a loading control. **(E)** Quantification of relative H3K27ac intensity in mouse brain sections from Gly7- and Gly73-NLS-expressing mice (n=3). **(F)** Quantification of relative H3K27ac intensity in the optogenetic polyG system under dark or blue-light conditions (n=3). **(G)** Representative images of H3K27ac immunostaining in the indicated groups. Scale bar, 5 μm. **(H)** Representative images of H3K27ac intensity in the optogenetic polyG system under dark or blue-light conditions. Scale bar, 10 μm. Data are means ± SEM and analyzed with an unpaired two-tailed Student’s t test (**C, E, F**) and one-way ANOVA (**B**). ‘‘ns’’ represents non-significant; *p < 0.05, and **p < 0.01.

Importantly, cells bearing visible intranuclear aggregates displayed lower H3K27ac signal than cells without aggregates (Supplemental Figure 6, A and B). Consistent with this, blue-light-induced formation of intranuclear opto-polyG aggregates significantly reduced H3K27ac intensity compared with the dark condition, indicating that polyG aggregation, rather than the soluble nuclear polyG, contributes to the reduced histone acetylation (Figure 5, F and H).

### Reduced H3K27 acetylation is observed in inclusion-bearing cells of human tissues

We next investigated H3K27 acetylation in human tissues. Brain autopsy of a genetically confirmed NIID patient, who was reported previously(31), showed that neurons containing intranuclear inclusions exhibited much lower H3K27ac signal than neighboring inclusion-negative neurons (Figure 6, A and B). Moreover, immunofluorescence analysis of biopsy samples, including skin, kidney, sural nerve and gallbladder, from patients with NIID, FXTAS, and OPML showed that cells containing intranuclear inclusions exhibited lower H3K27ac signal than neighboring inclusion-negative cells, suggesting a lack of tissue specificity (Figure 6, C-K). Together, these findings indicate that intranuclear inclusion is associated with reduced H3K27 acetylation across overexpressing models and human tissues of polyG diseases.

**Figure 6.**
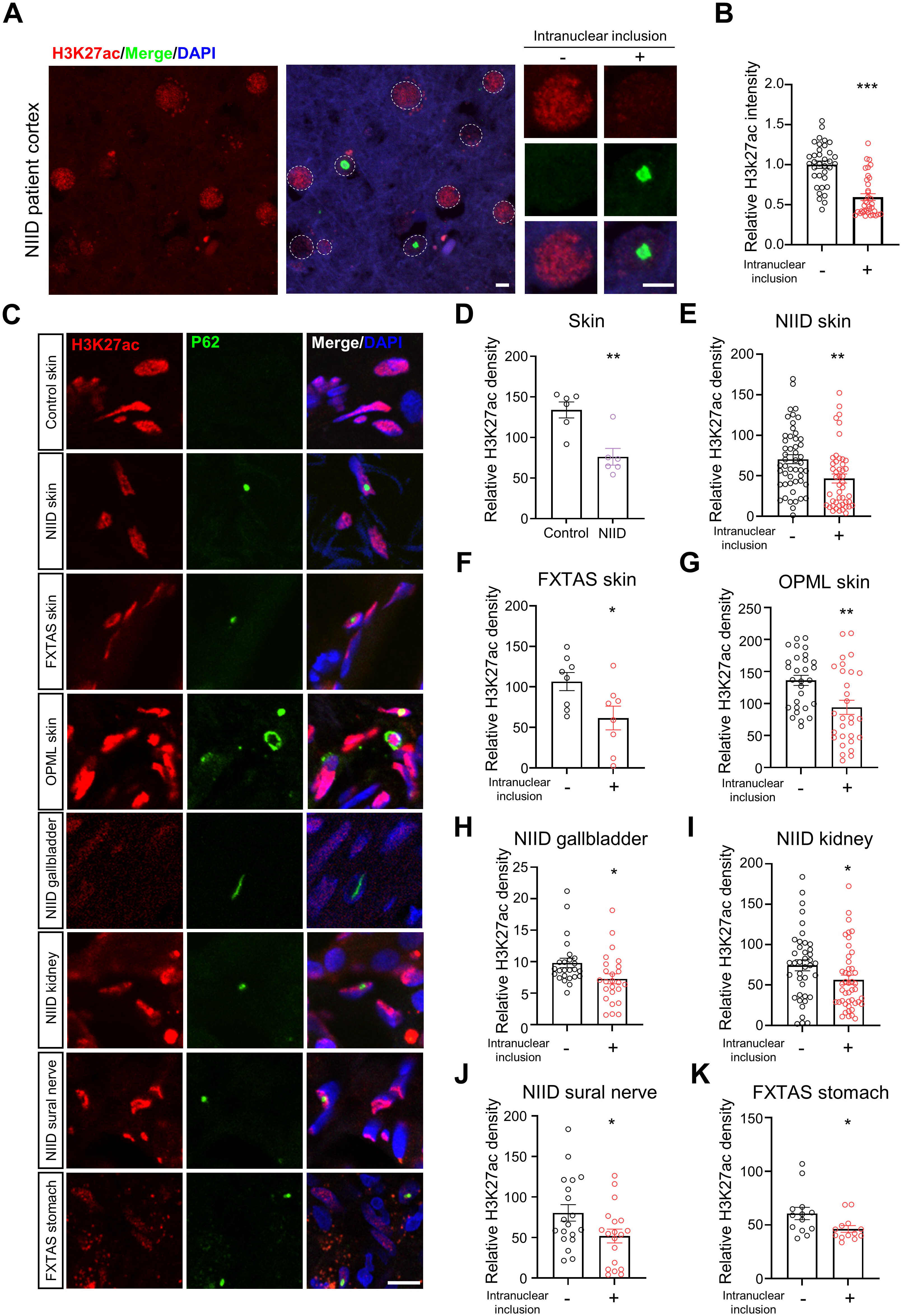
Inclusion-associated reduction of H3K27 acetylation in human polyG disease tissues. **(A)** Representative images of postmortem human brain sections stained for p62 and H3K27ac. Neurons containing intranuclear inclusions showed lower H3K27ac signal than neighboring inclusion-negative neurons. Scale bar, 5 μm. **(B)** Quantification of relative H3K27ac intensity in inclusion-positive and inclusion-negative neurons from human NIID brain tissue (30 neurons per group). **(C)** Representative immunofluorescence images of H3K27ac and p62 staining in skin, gallbladder, kidney, sural nerve, and stomach samples from control individuals and patients with NIID, FXTAS, or OPML. Scale bar, 10 μm. **(D)** Quantification of relative H3K27ac intensity in skin samples from control individuals and patients with NIID (n=6–7). **(E-K),** Quantification of relative H3K27ac intensity in inclusion-negative and inclusion-positive cells from NIID skin, FXTAS skin, OPML skin, NIID gallbladder, NIID kidney, NIID sural nerve, and FXTAS stomach. Data are means ± SEM and analyzed with an unpaired two-tailed Student’s test (**B, D-K**). *p < 0.05, **p < 0.01, and ***p < 0.001.

### HDAC3 upregulation is associated with intranuclear polyG-induced epigenetic repression

Given the concurrent reduction in chromatin accessibility and H3K27 acetylation, we next sought to identify candidate epigenetic regulators that might contribute to this repressive chromatin state. As histone acetylation is dynamically controlled by the opposing activities of histone deacetylases (HDACs) and histone acetyltransferases (HATs)(32), we examined their expression levels. Among the candidates analyzed, *Hdac3* was one of the few factors that showed a consistent increase in both the cortex and cerebellum of Gly73-NLS mice relative to controls (Supplementary Table 1 and Figure 7A). We therefore selected HDAC3 for further validation. Western blot analysis confirmed that HDAC3 protein abundance was elevated in brains from Gly73-NLS mice compared with controls (Figure 7, B and C). Consistently, immunofluorescence staining also showed increased HDAC3 intensity in the Gly73-NLS group (Supplemental Figure 6, C and D). Together, these findings identify HDAC3 upregulation as a molecular alteration associated with intranuclear polyG aggregation and raise the possibility that increased HDAC3 contributes to the reduction in H3K27 acetylation observed in this model.

**Figure 7.**
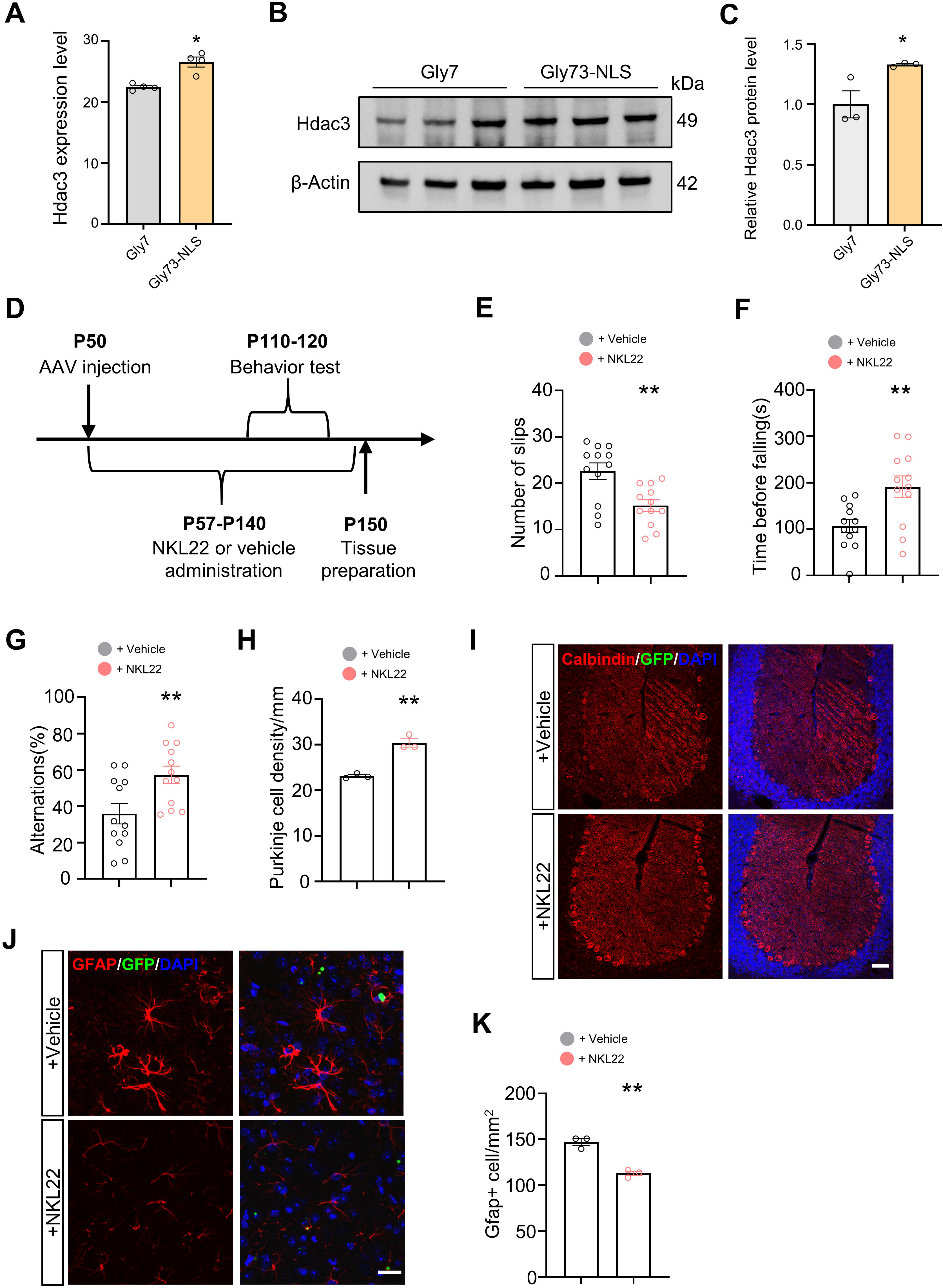
HDAC3 upregulation and partial rescue by HDAC inhibition in the Gly73-NLS mouse model. **(A)** RNA-seq-based expression level of *Hdac3* in mouse brains from the indicated groups, shown as FPKM values (n=4). **(B)** Representative immunoblot of HDAC3 in brain lysates from Gly7- and Gly73-NLS-expressing mice. β-actin served as a loading control. **(C)** Quantification of relative HDAC3 protein levels shown in (**B**) (n=3). **(D)** Schematic of the NKL22 treatment paradigm. Mice received AAV injection at P50, followed by subcutaneous administration of NKL22 or vehicle from P57 to P140. Behavioral analyses were performed between P110 and P120, and tissues were collected at P150. **(E)** Quantification of slips in the notched bar test in Gly73-NLS mice treated with vehicle or NKL22 (n=12). **(F)** Rotarod performance shown as latency to fall in Gly73-NLS mice treated with vehicle or NKL22 (n=12). **(G)** Y-maze spontaneous alternation in Gly73-NLS mice treated with vehicle or NKL22 (n=12). **(H)** Quantification of calbindin-positive Purkinje cell density (n=3). **(I)** Representative calbindin immunostaining images showing Purkinje cells in the cerebellum of vehicle- and NKL22-treated Gly73-NLS mice. Scale bar, 50 μm. **(J)** Representative GFAP immunostaining images in the cerebral cortex of vehicle- and NKL22-treated Gly73-NLS mice. Scale bar, 20 μm. **(K)** Quantification of GFAP-positive cell density (n=3). Data are means ± SEM and analyzed with an unpaired two-tailed Student’s t test (**A, C, E, F, G, H, K**). *p < 0.05 and **p < 0.01.

### HDAC3 inhibition ameliorates polyG-induced molecular, behavioral, and pathological deficits

To test whether the epigenetic alterations associated with intranuclear polyG aggregation are functionally relevant to the observed phenotype, we examined pharmacological inhibition with NKL22, an HDAC1/3 inhibitor previously reported to ameliorate phenotypes in Huntington’s disease models(33). In polyG-expressing cells, NKL22 treatment increased EU incorporation and H3K27ac intensity, supporting a partial rescue of nascent RNA synthesis and histone acetylation in vitro (Supplemental Figure 6, E-H).

We next evaluated the effect of NKL22 in vivo using the AAV-based Gly73-NLS mouse model. Mice received subcutaneous NKL22 treatment beginning 1 week after AAV injection, followed by behavioral testing and tissue analysis (Figure 7D). NKL22-treated mice showed significantly fewer slips in the notched-bar test, a longer latency to fall in the rotarod test, and increased spontaneous alternation in the Y-maze, indicating improvement in both motor and cognitive performance (Figure 7, E-G). However, NKL22 treatment did not significantly extend survival in Gly73-NLS mice (Supplemental Figure 6I). Histological analysis further showed that NKL22 attenuated neuropathological changes in the Gly73-NLS mice, including the Purkinje cell loss (Figure 7, H and I) and astrogliosis (Figure 7, J and K), whereas microglial activation was not significantly ameliorated (Supplemental Figure 6, J and M). In addition, NeuN staining and quantification revealed no significant difference in cortical neuronal density (Supplemental Figure 6, K and N), and the density of polyG aggregates remained unchanged (Supplemental Figure 6, I and O). Together, these findings suggest that pharmacological targeting of the HDAC pathway can partially ameliorate the molecular, behavioral, and pathological consequences of intranuclear polyG aggregation.

### Small polyG-containing peptides tend to form intranuclear aggregates at low expression levels

Findings of this study and our previous study suggest that the intranuclear localization of polyG aggregates was influenced by polyG length, the C-terminal sequence, engineered localization signals, and fusion tag size under overexpression conditions(11). However, these factors are still insufficient to explain the striking nuclear predominance seen in human patients, as naturally occurring polyG-containing peptides are small (∼10 kDa usually) and do not contain canonical NLS. Because the nucleus has a more limited capacity than the cytoplasm to clear protein aggregates(34, 35), and polyG proteins are expressed at substantially lower levels than polyQ proteins (Supplementary Table 2 and Supplemental Figure 7A), we reasoned that the low expression levels and long-term accumulation of polyG-containing peptides may facilitate the formation of intranuclear aggregates. To test this hypothesis, we performed an independent experiment using our previously established AAV expressing HA-tagged uN2CpolyG, a more native polyG with a molecular weight of 11.9 kDa(18). Following low- or high-dose injection into C57BL/6 mice, the low-dose group exhibited a significantly higher proportion of intranuclear aggregates in the cortex than the high-dose group at 4 months after injection (Supplemental Figure 7, B-D).

## DISCUSSION

In this study, we demonstrate that intranuclear polyG aggregates are substantially more neurotoxic than cytoplasmic aggregates in both cellular and mouse models. We further show that nuclear polyG aggregation is associated with a repressive epigenetic state characterized by impaired nascent RNA synthesis, reduced chromatin accessibility, decreased H3K27 acetylation, and increased HDAC3 expression, with validation in human tissues. The partial rescue achieved by pharmacological HDAC inhibition supports the functional relevance of this pathway in polyG-induced neurodegeneration.

A long-standing enigma of polyG diseases is the predominant intranuclear localization of inclusions observed in patients. Unlike intrinsically nuclear-targeted DPRs such as polyPR in ALS/FTD(36), our findings suggest that polyG tracts themselves are not preferentially directed to a specific subcellular compartment. The abundant cytoplasmic polyG inclusions commonly observed in widely used cellular and animal models may therefore represent an artifact of overexpression. In contrast, slow, low-level expression of polyG-containing peptides favors intranuclear aggregation, more closely mirroring the pathology observed in human disease. However, toxicity under these low-expression conditions is minimal and insufficient to recapitulate key disease phenotypes within the lifespan of cultured cells or mice. To address this limitation, we employed overexpression constructs bearing tandem 2×EGFP tags, which enabled improved control of nuclear localization. Although this design increases the molecular weight of the polyG proteins, the resulting phenotypic, pathological, and transcriptional features closely match those observed in our previous uN2CpolyG overexpression study(18). Together, these findings support the use of 2×EGFP-tagged constructs as a practical and reliable approach for dissecting polyG toxicity mechanisms.

Using this system, we found that the pathogenicity of polyG proteins is strongly associated with their intranuclear localization and aggregation. Despite showing the lowest mRNA and protein abundance among the polyG-expressing groups, the nuclear-targeted Gly73-NLS construct elicited the most severe pathological and behavioral phenotypes. Importantly, phenotypic severity followed a clear gradient, with Gly73-NLS mice most affected, Gly73 mice intermediate, Gly73-NES mice least affected, and controls largely normal. This pattern suggests that neurodegenerative burden scales with the proportion of polyG aggregates localized to the nuclei, highlighting that intranuclear accumulation is not simply associated with toxicity, but is a critical driver of it.

The nucleus plays a central role in highly organized processes essential for neuronal survival, including transcription, chromatin maintenance, RNA processing, and nucleocytoplasmic communication(37, 38). At the same time, the nucleus represents a vulnerable compartment for aberrant protein aggregation in multiple neurodegenerative diseases(39, 40). Beyond polyG diseases, intranuclear inclusions have been detected in some other repeat expansion disorders, including polyQ diseases, polyalanine (polyA) diseases, and *C9ORF72*-associated ALS/FTD(21). Notably, prior studies in Huntington’s disease and Kennedy’s disease mouse models demonstrated that intranuclear polyQ can exert greater toxicity than cytoplasmic polyQ despite identical protein composition(41–43), raising the possibility that selective nuclear vulnerability may be a convergent pathogenic mechanism for repeat expansion disorders harboring intranuclear inclusions.

Despite these observations, the molecular basis of nuclear vulnerability remains poorly understood. Potential contributing factors may include limited nuclear protein quality control capacity, impaired nucleocytoplasmic transport, altered chromatin organization, and sequestration of transcriptional regulators.

In this study, we focus on the transcriptional and epigenetic dysregulation related to intranuclear polyG aggregates. Although widespread transcriptional dysregulation is an important pathogenic feature of several repeat expansion disorders(44, 45), its role in polyG diseases remains largely unexplored. Our data demonstrate that intranuclear polyG aggregation results in impaired nascent RNA synthesis both in vitro and in vivo. This can cause multiple downstream abnormalities and help contextualize previous findings that polyG pathology is associated with nucleolar stress, impaired ribosome biogenesis, and translational inhibition(46). Impaired nascent RNA synthesis may lie upstream of at least part of these abnormalities, thereby contributing to broader defects in RNA homeostasis, ribosome function, and protein synthesis.

Transcriptional output is regulated by cellular metabolism and epigenetic modifications(47). Our data further suggest that the observed transcriptional impairment is accompanied by broader epigenetic remodeling. In particular, intranuclear polyG aggregation is associated with globally reduced chromatin accessibility, indicating that its effects extend beyond selected genes or pathways to a widespread loss of transcription-permissive chromatin states. This global chromatin closing may underlie the reduced nascent RNA synthesis observed in our models by broadly limiting access of transcription factors and RNA polymerase II-associated machinery to genomic DNA. More broadly, these findings raise the possibility that intranuclear polyG aggregates remodel the nuclear landscape toward a state less compatible with transcriptional homeostasis, which is increasingly recognized as an important feature of neurodegenerative diseases(48, 49).

The reduced H3K27ac, observed in both Gly73-NLS-overexpressing models and patient tissues across various polyG diseases, further supports a repressive chromatin environment associated with intranuclear polyG aggregation. We focused on H3K27ac because it is a well-established marker of transcriptionally accessible chromatin, particularly at active regulatory elements, and thus an informative readout of transcriptional competence(30, 50). Although other histone acetylation markers may also be altered, H3K27ac provided a relevant entry point for assessing whether nuclear polyG aggregation is associated with a less permissive epigenetic landscape(51). Taken together, the concomitant loss of chromatin accessibility and H3K27ac provides a mechanistic framework linking intranuclear polyG aggregation to reduced transcriptional output.

It remains unclear how intranuclear polyG aggregation causes reduced H3K27ac. The lack of disease and tissue specificity observed in human samples supports an upstream and conservative pathway repressing H3K27ac. The correlation between the intranuclear aggregation burden and the extent of reduction in nascent RNA synthesis and H3K27ac suggests a potential role of intranuclear macromolecular crowding caused by polyG aggregates, which can lead to an excluded volume effect and an environment unfavoring chromatin accession and transcription activity(52, 53). In addition, whether and how disrupted phase separation, which has been revealed in polyG diseases(54), contribute to such an epigenetic repression deserve further investigation.

Our observation of increased HDAC3 provides a mechanistic rationale for therapeutic intervention. Consistent with prior studies implicating HDAC3 in transcriptional repression, chromatin dysregulation, and neuronal vulnerability in polyQ diseases, we selected NKL22 as a mechanism-informed pharmacological tool(55–57). In our model, NKL22 partially restored H3K27ac levels and nascent RNA synthesis and ameliorated behavioral and pathological abnormalities, supporting a functional contribution of HDAC-dependent epigenetic dysregulation to polyG diseases. However, the rescue was incomplete, with the absence of a survival benefit. Such a dissociation between phenotypic improvement and survival has also been reported for NKL22 in Huntington’s disease models(33, 58), suggesting that HDAC3-associated dysregulation is important but not exclusive in polyG toxicity.

Overall, this study provides evidence that the pathogenicity of polyG proteins is strongly shaped by their subcellular localization and aggregation, with intranuclear inclusions representing the more toxic species. Our findings further suggest that a major downstream consequence of intranuclear polyG aggregation is a transcriptionally repressive chromatin environment, characterized by reduced chromatin accessibility, diminished H3K27ac, and impaired nascent RNA synthesis, potentially mediated in part by HDAC3 dysregulation. The partial rescue achieved by pharmacological HDAC inhibition further supports the functional importance of this pathway. These results not only advance the mechanistic understanding of polyG diseases and related repeat expansion disorders harboring intranuclear inclusions, but also point to epigenetic modulation as a promising therapeutic direction.

## METHODS

### Sex as biological variable

Sex was not considered as a biological variable.

### Plasmid constructs

The AAV packaging constructs were derived from our previously established pAAV-CAG backbone. To optimize expression, a shortened CAG promoter was generated, resulting in the pAAV-sCAG vector. The GGN11 and GGN73 repeat sequences were amplified from our previously constructed plasmid pAAV-ATG-GGN100dCT. These sequences were fused in-frame with either 0, 1, or 2 copies of GFP, followed by a C-terminal 3×FLAG tag. To manipulate subcellular localization, the constructs were further engineered to contain either two tandem nuclear localization signals (2×NLS) or a nuclear export signal (NES). The resulting expression cassettes were subsequently cloned into the pAAV-sCAG vector.

For optogenetic induction of polyG aggregation, a Gly49-mCherry-CRY2clust-NLS overexpression plasmid was constructed using pCAG-GFP (Addgene plasmid #16664) as the backbone. The GGN49 repeat sequence was amplified from our previously constructed plasmid pAAV-ATG-GGN100dCT. The mCherry-CRY2clust module was synthesized on the basis of a previously reported optogenetic clustering system, in which the CRY2clust sequence was derived from the C-terminal extension of CRY2PHR and fused to mCherry(26). The GGN49 fragment was then inserted in-frame upstream of CRY2clust-mCherry-NLS to generate the final Gly49-mCherry-CRY2clust-NLS construct. All recombinant plasmids were transformed into Escherichia coli (F-DH5α Stable strain, Shanghai Weidi Biotechnology) and cultured at 37 °C. All recombinant plasmids were verified by Sanger sequencing.

### Cell culture and transfection

SH-SY5Y cells (ATCC, CRL-2266) were cultured in DMEM/F12 (high glucose) supplemented with 10% fetal bovine serum (FBS, Sigma, F0850) and 100 IU/mL penicillin–streptomycin (Gibco, 15140163). Cells were cultured at 37 °C in a humidified incubator with 5% CO . SH-SY5Y cells were seeded into 6-well, 12-well, or 24-well plates. Approximately 18 h after seeding, cells were transfected with plasmid DNA using Lipo8000 transfection reagent (Beyotime Biotechnology) according to the manufacturer’s instructions. The culture medium was replaced 8–12 h post-transfection. Cells were harvested 36–48 h after transfection for subsequent experiments.

### Cell death and viability assay

Cell viability of SH-SY5Y cells was evaluated by flow cytometry using Annexin V staining. Briefly, cells were seeded into 6-well plates and transfected with the indicated plasmids. At the designated time points, culture supernatants were collected, and adherent cells were detached using trypsin. The detached cells were combined with the corresponding supernatants and centrifuged at 500 × g at 4 °C. The cell pellets were washed with Hank’s Balanced Salt Solution (HBSS), gently resuspended, and incubated with Annexin V-APC (KGA1105-20, KeyGEN BioTECH) for 10 min in the dark according to the manufacturer’s instructions. Flow cytometric analysis was performed using a MA900 Flow Cell Sorter (SONY), and at least 50,000 events were acquired per sample.

Cell death was further assessed by measuring lactate dehydrogenase (LDH) release using an LDH Cytotoxicity Assay Kit (C0017, Beyotime). SH-SY5Y cells were seeded into 24-well plates and transfected with the indicated plasmids. Culture supernatants were collected and analyzed for LDH activity according to the manufacturer’s protocol.

### Western blotting

Cells were lysed in RIPA lysis buffer supplemented with PMSF and protease/phosphatase inhibitor cocktails and incubated on ice for 10 min. Lysates were clarified by centrifugation at 12,000 × g for 10 min at 4 °C, and the protein concentration of the supernatants was determined using the Bradford assay. Equal amounts of protein were denatured by boiling and separated on 4–12% Tris–glycine SDS–PAGE gels, followed by transfer onto PVDF membranes. Membranes were blocked with 5% non-fat dry milk in TBST for 1 h at room temperature and then incubated overnight at 4 °C with the following primary antibodies: mouse monoclonal anti-DYKDDDDK tag (1:2000; 66008-4-Ig, Proteintech), rabbit polyclonal anti-HDAC3 (1:2000; A2139, Abclonal), rabbit polyclonal anti-H3K27ac (1:2000, ab4729, Abcam), or rabbit monoclonal anti-β-Actin (1:100,000; AC026, Abclonal). After washing with TBST, membranes were incubated with horseradish peroxidase-conjugated goat secondary antibodies (1:2000; Beyotime) for 1 h at room temperature. Protein bands were visualized using enhanced chemiluminescence (ECL) reagents and imaged with a VersaDoc imaging system (Bio-Rad).

### Real-time qPCR

Total RNA was extracted from cultured cells and reverse-transcribed into cDNA using the HiScript II Q RT SuperMix kit (R222, Vazyme) according to the manufacturer’s instructions. The resulting cDNA was used as a template for quantitative PCR amplification with SYBR qPCR Master Mix (Q711-00, Vazyme). Quantitative real-time PCR was performed on an ABI QuantStudio™ 3 Real-Time PCR System (Thermo Fisher Scientific). Relative mRNA expression levels were calculated using the comparative Ct (2^−ΔΔCt) method and normalized to the indicated internal control gene.

The following primer pairs were used for qPCR:

WPRE-F: 5’-GGATACGCTGCTTTAATGCCTTT-3’ WPRE-R: 5’-GCGAAAGTCCCGGAAAGGAG-3’ HDAC3-F(human): 5’-AGGCCTCCCAACATGACATG-3’ HDAC3-R(human): 5’-TGTGTAACGCGAGCAGAACT-3’ ACTB-F (human): 5’-GGCTGTATTCCCCTCCATCG-3’ ACTB-R (human): 5’-CCAGTTGGTAACAATGCCATGT-3’ Actb-F (mouse): 5’-TGTTACCAACTGGGACGACA-3’ Actb-R (mouse): 5’-GGGGTGTTGAAGGTCTCAAA-3’

### Animals

All animal procedures were approved by the Ethics Committee of Zhongshan Hospital, Fudan University (Approval ID: 2025-221), and were conducted in accordance with institutional guidelines for animal care and use. Male C57BL/6 wild-type mice (mean body weight: 22 g; Shanghai JieSiJie Laboratory Animal Co., Ltd.) received intravenous injection of AAV at P50. Following injection, mice were maintained for 4–8 months in a temperature-controlled facility under a 12-h light/dark cycle with ad libitum access to food and water. For molecular analyses, a subset of mice was sacrificed by cervical dislocation for brain dissection and subsequent sequencing experiments. For histological analysis, mice were deeply anesthetized with sodium pentobarbital (50 mg/kg, intraperitoneally) and transcardially perfused with ice-cold phosphate-buffered saline (PBS, pH 7.4), followed by 4% paraformaldehyde (PFA) in PBS. Brain tissues were collected at P150. The remaining mice were monitored for survival analysis and maintained until P400. Body weight and survival status were recorded regularly throughout the experimental period.

### AAV production and intravenous injection

Recombinant AAV2/PHP.eB vectors were packaged by OBiO Technology Inc. (Shanghai, China). Viral particles were generated by triple transfection of HEK293 cells with the pAAV-sCAG expression constructs (Gly7, Gly73, Gly73-NLS, or Gly73-NES), together with the helper plasmids pUCmini-iCAP-PHP.eB and pADDELTA-F6. Recombinant AAV vectors were purified from cell lysates using double cesium chloride gradient ultracentrifugation, followed by dialysis and concentration in sterile PBS. Viral genome titers were determined by quantitative real-time PCR and expressed as viral genomes per milliliter (vg/mL). For in vivo experiments, each mouse received a single intravenous injection of 200 μL sterile Hank’s Balanced Salt Solution (HBSS) containing 3 × 10¹¹ vg of AAV2/PHP.eB particles.

### Behavioral tests

Rotarod test: Motor coordination and balance were assessed using a rotating rod apparatus (47650 Rota-Rod, UGO Basile). The rotation speed was gradually accelerated from 4 to 40 rpm over 5 min. Each mouse underwent three trials with inter-trial intervals of 1–1.5 h. The latency to fall from the rod was recorded for each trial, and the average latency was used as an index of motor performance.

Notched bar test: Motor coordination and balance were further evaluated using a notched wooden bar as previously described(18). The apparatus consisted of a 2-cm-wide and 50-cm-long wooden bar with 12 square platforms (2 cm each) separated by 13 gaps (2 cm each), and two rectangular platforms at both ends. The bar was elevated 25 cm above the desktop. Mice were trained to cross the bar twice before testing and then tested in three consecutive trials. Each instance in which a hind paw slipped into a gap was recorded as one slip. The total number of slips across the three trials was calculated as the final score. If a mouse fell from the bar, it was scored as 12 slips.

Y maze: Working memory was evaluated using a Y-maze apparatus with three identical arms (30 × 6 × 15 cm). Each mouse was placed at the end of one arm and allowed to explore freely for 10 min. Arm entries were recorded, and spontaneous alternation behavior was analyzed using an automated tracking system. Alternation was defined as consecutive entries into three different arms. The alternation rate was calculated as the percentage of actual alternations relative to the maximum possible alternations. A lower alternation rate was considered indicative of impaired working memory.

Novel object recognition test: Mice were placed in the center of an open field arena (40 × 40 × 40 cm). During the habituation phase, mice were placed individually in the empty arena and allowed to explore freely for 5 min. Twenty-four hours later, during the training phase, two identical objects were placed in the arena, and mice were allowed to explore for 5 min. After another 24 h, one of the familiar objects was replaced with a novel object of similar size but different shape. Mice were then allowed to explore for 5 min. Exploration was defined as directing the nose toward or touching the object. The time spent exploring the novel and familiar objects was recorded and compared among groups.

### Immunofluorescent staining

PFA-fixed mouse brains were cryoprotected in 30% sucrose at 4 °C for at least 72 h, frozen in the embedding medium and serially cryo-sectioned at 16 µm for histological analysis. For immunofluorescence staining, sections were permeabilized in TBS with 0.5% Triton X-100, and blocked for 2 h in TBS with 5% donkey serum (S2170-500, Biowest). Floating sections were incubated overnight at 4°C in primary antibodies from different species diluted in blocking buffer and then in secondary antibodies sequentially. Secondary antibodies conjugated with Alexa Fluor dyes (1:1000, all from Jackson ImmunoResearch) were applied for 1 hour at room temperature, followed by rinsing in TBS, counterstaining with 4,6-diamidino-2-phenylindole (DAPI) (200 ng/mL, Sigma) for 5 minutes, and mounting. Omission of primary antibodies eliminated the staining signals. The following primary antibodies were used: rabbit anti-GFP (1:1000; 66002-1-Ig, Proteintech), rabbit anti-p62/SQSTM1 (1:200; 18420-1-AP, Proteintech), rabbit anti-Gfap (1:2000; Z0334, Dako), rabbit anti-Calbidin-D28K (1:500; CB-38a, Swant), goat anti-Iba1 (1:500; 019-19741, Wako), rabbit anti-DYKDDDDK tag (1:2000; 66008-4-Ig, Proteintech), rabbit anti-HDAC3 (1:2000; A2139, Abclonal), rabbit anti-H3K27ac (1:2000, ab4729, Abcam), and rabbit anti-NeuN (1:1000; ab177487, Abcam).

### Transmission electron microscopy

For transmission electron microscopy (TEM) analysis, mouse tissue samples were fixed in 2.5% glutaraldehyde, followed by post-fixation in 1% osmium tetroxide. Samples were then dehydrated through a graded ethanol series and embedded in epoxy resin. Ultrathin sections were prepared and stained with uranyl acetate and lead citrate. Images were acquired using either a Philips CM120 transmission electron microscope operated at 60 kV or a Tecnai G2 Spirit TWIN transmission electron microscope operated at 80 kV.

### EU staining

Cells were incubated with 1 mM 5-ethynyl uridine (EU) at 37 °C for 2 h at 48 h post-transfection. Following EU labeling, culture medium was removed and cells were fixed with 4% PFA for 15 min at room temperature. After washing with TBS, cells were permeabilized with TBS with 0.5% Triton X-100 for 15 min at room temperature. Subsequently, a Click reaction buffer was freshly prepared according to the manufacturer’s instructions and added to the cells for 30 min at room temperature in the dark. After completion of the Click reaction, nuclei were counterstained for imaging.

Collected brains were embedded in Tissue-Tek® O.C.T. Compound (Sakura) and rapidly frozen on dry ice. Frozen tissues were sectioned at 14 μm thickness using a cryostat. Sections were fixed in 4% PFA in PBS for 10 min at room temperature, followed by washing with TBS. Tissue sections were permeabilized with 0.5% Triton X-100 in TBS for 30 min at room temperature. EU incorporation was detected using the BeyoClick™ EU RNA Synthesis Kit with AF555 (R0305S, Beyotime) according to the manufacturer’s instructions. For combined EU Click chemistry and immunohistochemistry, sections were briefly re-fixed in 4% PFA for 5 min at room temperature following the Click reaction. Sections were then washed with TBS and blocked for 2 h at room temperature in TBS containing 5% donkey serum (S2170-500, Biowest). Primary antibody incubation was performed overnight at 4 °C using rabbit anti-NeuN (1:1000; ab177487, Abcam).

### RNA-seq

Bulk RNA-seq was performed on cerebral cortical tissue collected from mice in the Gly7, Gly73, Gly73-NLS, and Gly73-NES groups at P150 (n = 4 per group). Total RNA was extracted using TRIzol reagent (Invitrogen). Library preparation and paired-end sequencing (2 × 150 bp) were performed on an Illumina NovaSeq 6000 platform by GENEWIZ Biotechnology (Suzhou, China).

Sequencing data quality was assessed by FastQC software (v0.10.1), and raw reads were filtered by Cutadapt (version 1.9.1). Reads were mapped to the *Mus musculus* reference genome (GRCm39), and expression levels were estimated by FPKM. Differential expression analysis between Gly7 and the other groups was performed using the DESeq2 R package (V1.26.0). P values were adjusted using Benjamini and Hochberg’s approach for controlling the false discovery rate (FDR). Genes with an adjusted P value<0.05 and | log2 (fold change) |> 1.0 found by DESeq2 were assigned as differentially expressed. Functional enrichment analysis was performed using Metascape. Gene Ontology Biological Process enrichment was conducted on the upregulated genes identified in each comparison group relative to Gly7.

### ATAC-seq

Cells were collected and counted at room temperature using the Countstar Rigel S2(Shanghai Ruiyu, FL20447, CHINA). The requisite number of cells for the experiment was lysed for 5 minutes to extract the nuclei. The nuclei were then suspended in a transposition reaction system containing Tn5 transposase. This was followed by incubation and DNA purification. The resulting product was subjected to PCR amplification with the introduction of specific indexes. The PCR amplification conditions were as follows: incubation at 72°C for 3 minutes, pre-denaturation at 98 ° C for 1 minute, denaturation at 98 ° C for 10 seconds, annealing at 60°C for 25 seconds, and extension at 72°C for 25 seconds, for a total of 13 to 15 cycles, with a final extension at 72°C for 5 minutes.

Library fragments of approximately 200-700 bp were obtained through bead selection. The concentration of the library was measured using Qubit (Thermo, Qubit3.0, USA) and the integrity of the fragments was assessed using a Bioanalyzer 2100(Agilent, CA, USA) Perform PE150 sequencing using the Illumina NovaSeqXP according to standard protocols.

Raw sequencing reads were processed to remove adapter sequences and low-quality bases, and clean reads were aligned to the reference genome using Bowtie2. Duplicate reads, low-quality alignments, and mitochondrial reads were removed before downstream analysis. Peaks were called using MACS2 and annotated using ChIPseeker. Differential accessibility analysis between groups was performed using DiffBind, and functional enrichment analysis of genes associated with differential peaks was carried out using GO and KEGG analyses. Biological replicates were included for each group, as indicated in the corresponding figure legends.

### Opto-stimulation

Cells expressing opto-constructs were stimulated with a 488 nm laser on a custom-built LED blue-light array housed in a humidified incubator maintained at 37℃ with 5% CO2(59). Cells were protected from light between experiments and during fixation.

### Human samples

All human tissues were obtained and distributed under oversight by the Ethics Committee of Zhongshan Hospital, Fudan University (Approval ID: B2024-478R; B2025-132). Brain tissue from a NIID patient was provided by the Shanghai Brain Bank, Fudan University. Skin tissues were obtained by punch or ordinary biopsy 10 cm above the lateral malleolus. All tissues were fixed in 4% PFA, cryoprotected in 30% sucrose and sectioned into 8-μm frozen sections. Written informed consent for publication was obtained where applicable.

### Microscopy

Fluorescent images were captured with an Olympus FV3000 confocal microscope using ×40 or ×60 objectives or an Olympus VS120 slide scanning system using a ×10 or ×20 objective.

### Quantification and statistical analysis

Data were analyzed and plotted using GraphPad Prism 9.0 software. Results are presented as mean ± SEM. Statistical significance was determined using Student’s t test, one-way ANOVA with Tukey’s or Dunnett’s multiple comparisons post-test, or Kruskal–Wallis test with Dunnett’s multiple comparisons post-test. P value less than 0.05 was considered statistically significant. The sample sizes and other details of statistical analyses are described in the figure legends.

## Supporting information

Supplementary mateiral

## Data availability

All data and resources that support the findings of this study are available from the corresponding author upon reasonable request.

## Acknowledgements

This work was supported by project grants from National Natural Science Foundation of China (Grant NO.82271499; 82301603), China Postdoctoral Science Foundation (Grant NO.2023TQ0078; 2023M730676), and Zhongshan Hospital Fudan University (Grant NO. 2024ZYYS-017). We thank Prof. Yang Zhengang, Prof. Li Wensheng, Dr. Gao Hongyang, Dr. Yu Zhang (Fudan University), Dr. Li Xiaoya (Shanghai Jiao Tong University), Dr. Guo Liang (Tsinghua University), and Dr. Chen Xiufei (Ningbo University) for their technical support and academic suggestions. We are deeply grateful to the patient who donated the brain tissue for this study.

## Author contributions

S.P.Z. and Y.Y.L. contributed to the conception and design of the study; S.P.Z., Y.Y.L., J.X.H., J.Z.L., L.Y.H. and Y.Z.L. contributed to the acquisition of the data; Y.Y.L. and J.X.H. contributed to the analysis of the data; Y.Y.L. and S.P.Z. contributed to the statistics; Y.Y.L. and S.P.Z. contributed to drafting the text and preparing the figures; J.D. and S.P.Z. supervised the study and revised the manuscript.

