## Supplementary mateiral for "Intranuclear polyglycine aggregation drives neurodegeneration through epigenetic repression of chromatin accessibility and transcription"

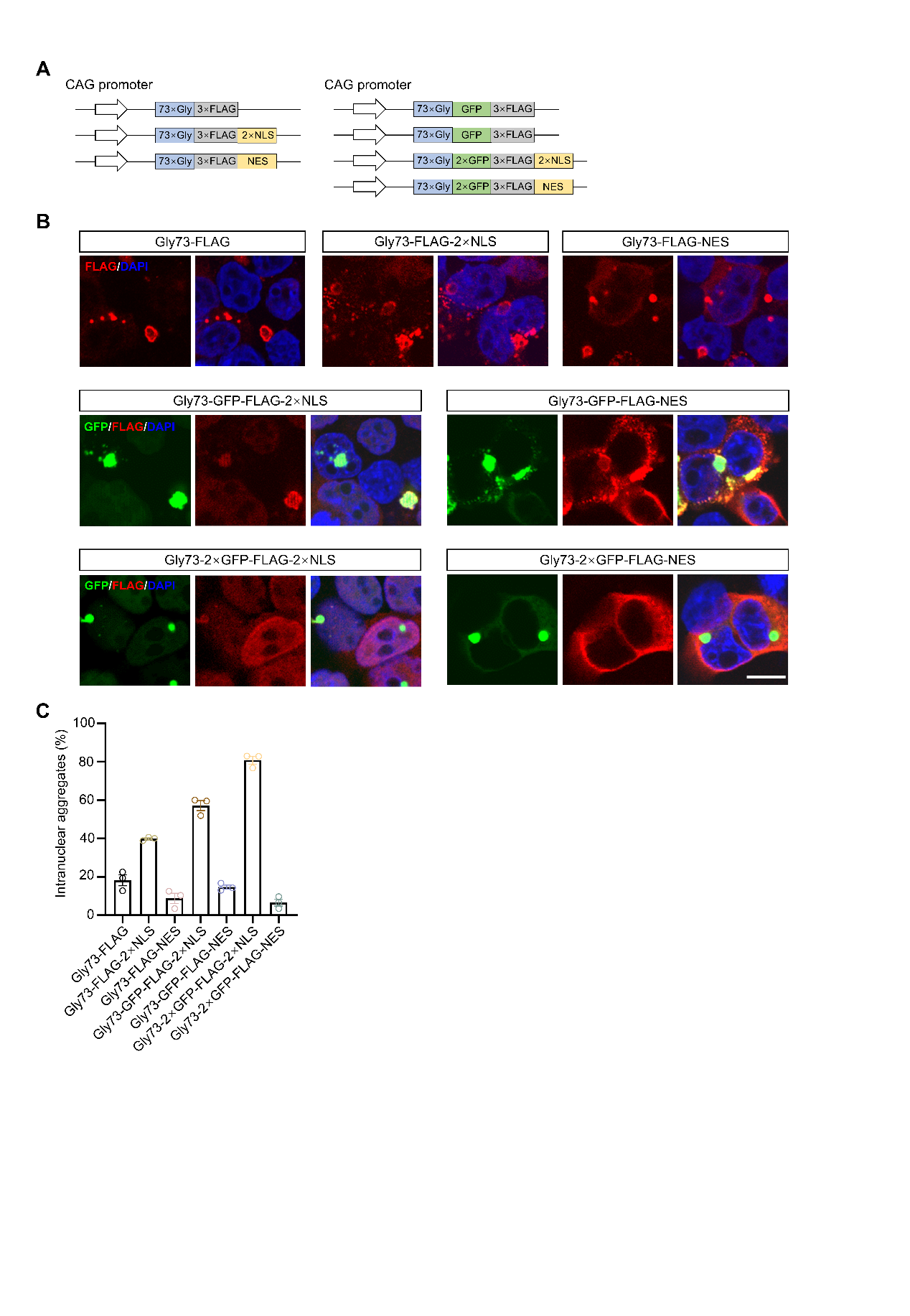
**Supplementary Materials**

**Supplemental Figure 1. Increasing fusion protein size improves control of polyG aggregate localization. (A)** Schematic of the indicated polyG constructs with different tagging strategies. Gly73 was fused to FLAG alone, EGFP-FLAG, or 2×EGFP-FLAG, with or without a 2×NLS or NES, to examine how protein size influences the subcellular distribution of polyG aggregates. **(B)** Representative fluorescence images of SH-SY5Y cells expressing the indicated constructs. Increasing fusion protein size progressively enhanced the ability of the localization signals to bias aggregates distribution, with the 2×EGFP/2×NLS construct showing the strongest enrichment of intranuclear aggregates. Scale bar, 10 μm. **(C)** Quantification of the percentage of intranuclear inclusions among total inclusions formed by the indicated constructs (n=3). Data are means ± SEM and analyzed with one-way ANOVA (**C**). ‘‘ns’’ represents non-significant; *p < 0.05, **p < 0.01, and ***p < 0.001.

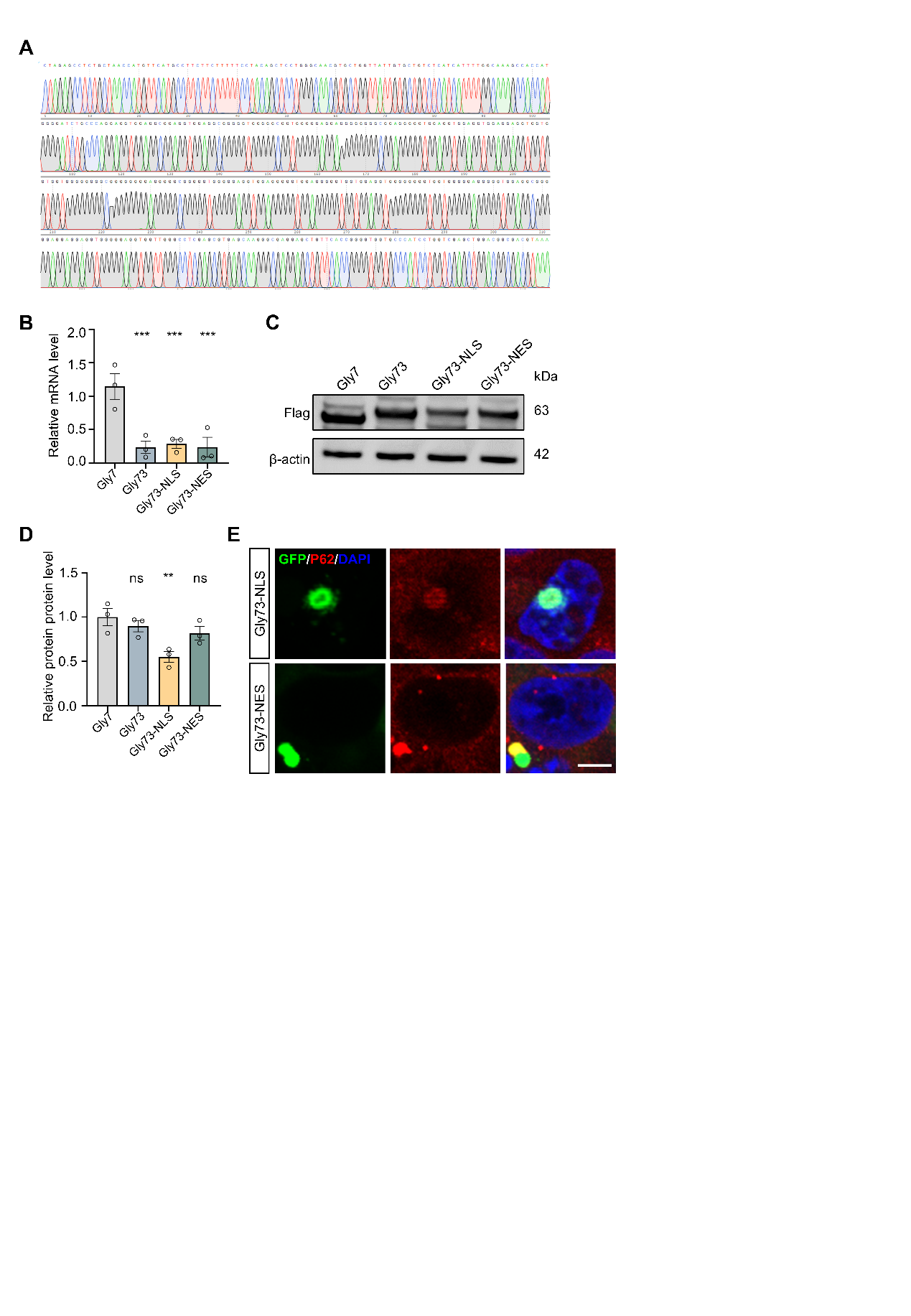
**Supplemental Figure 2. Validation of polyG reporter constructs and pathological characterization of polyG aggregates.** **(A)** Sanger sequencing traces confirming the repeat composition of the indicated polyG constructs. **(B)** qPCR analysis of the relative mRNA levels of Gly7, Gly73, Gly73-NLS, and Gly73-NES in transfected cells (n=3). **(C)** Representative immunoblot showing protein expression of the indicated constructs detected by anti-FLAG antibody. β-actin was used as a loading control. **(D)** Quantification of relative protein levels shown in (C) (n=3). **(E)** Representative confocal images of SH-SY5Y cells expressing Gly73-NLS or Gly73-NES, co-immunostained for GFP and p62. PolyG aggregates co-localized with p62. Scale bar, 5 μm. Data are means ± SEM and analyzed with one-way ANOVA (**B, D**). ‘‘ns’’ represents non-significant; **p < 0.01, and ***p < 0.001.

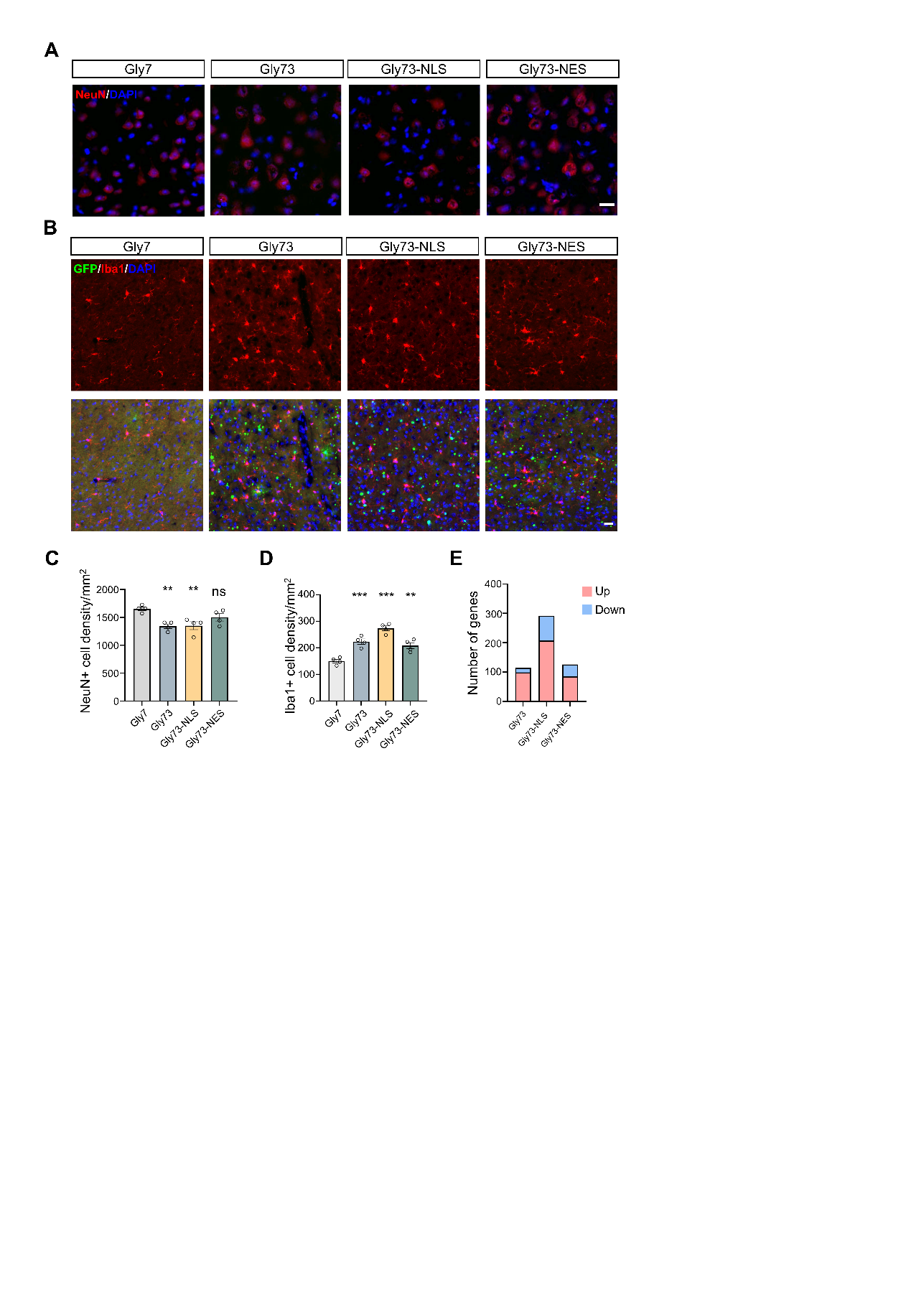
**Supplemental Figure 3. Neuronal loss and microglial activation in mice with nuclear and cytoplasmic polyG aggregation.** **(A)** Representative NeuN immunostaining images in the cerebral cortex of mice of the indicated groups at P150. Scale bar, 20 μm. **(B)** Representative Iba1 immunostaining images in the cerebral cortex of mice of the indicated groups at P150. Scale bar, 100 μm. **(C)** Quantification of NeuN-positive cell density in the cerebral cortex (n=4). **(D)** Quantification of Iba1-positive cell density (n=4). **(E)** Numbers of differentially expressed genes in the cerebral cortex of Gly73, Gly73-NLS, and Gly73-NES mice relative to Gly7 controls at P150. Data are means ± SEM and analyzed with a one-way ANOVA (**C, D**). ‘‘ns’’ represents non-significant; *p < 0.05, **p < 0.01, and ***p < 0.001.

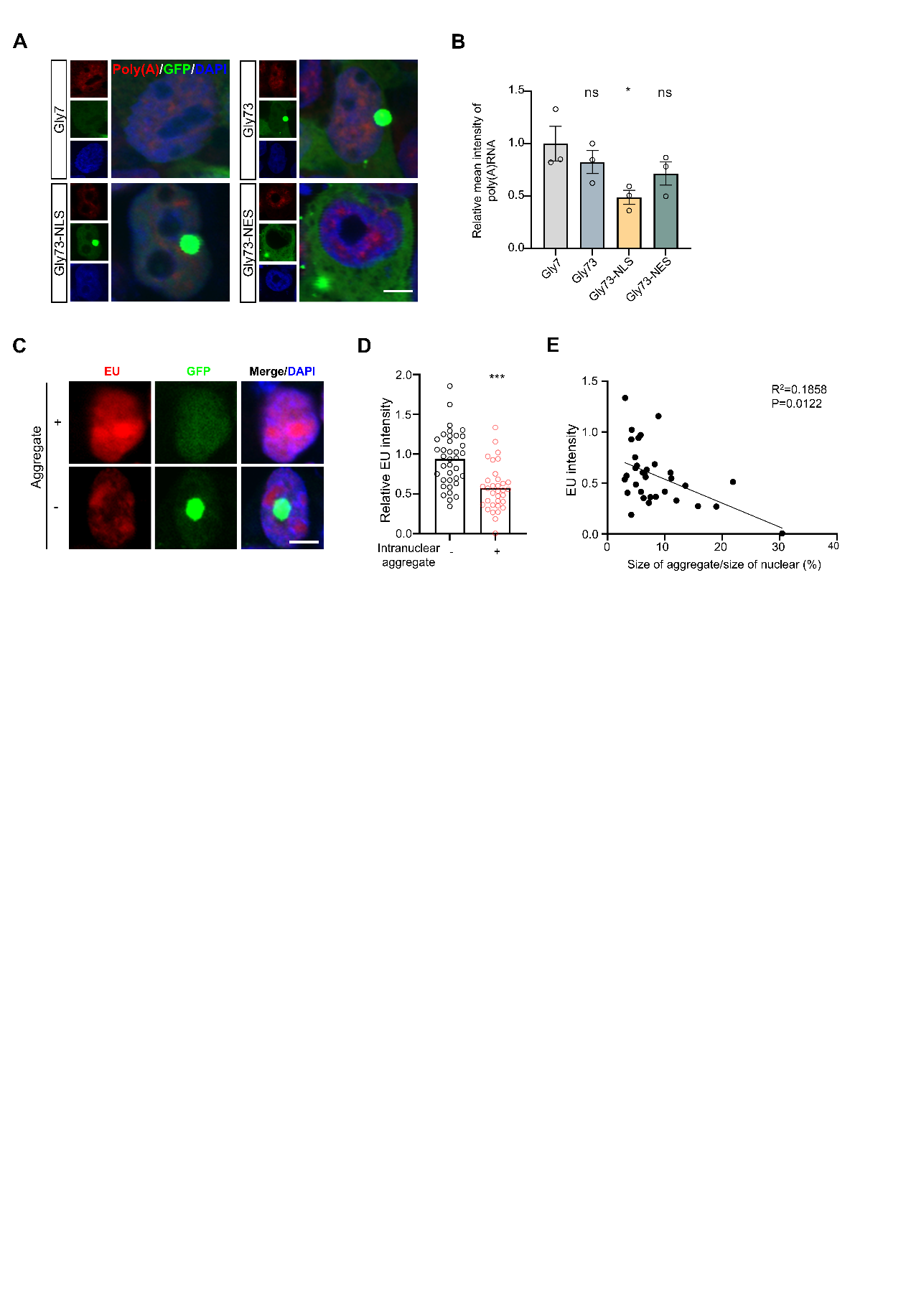
**Supplemental Figure 4. Intranuclear polyG aggregation is associated with aggregate-dependent suppression of nascent RNA synthesis and reduced mRNA levels.** **(A)** Representative images of oligo(dT) staining in cells expressing Gly7, Gly73, Gly73-NLS, or Gly73-NES. Scale bar, 5 μm. **(B)** Quantification of relative mean intensity of mRNA in the indicated groups (n=3). **(C)** Representative images of EU labeling in aggregate-negative and aggregate-positive cells expressing Gly73-NLS. Scale bar, 5 μm. **(D)** Quantification of relative EU intensity in aggregate-negative and aggregate-positive cells (10 cells from each of three independent experiments per group). **(E)** Correlation analysis between EU intensity and aggregate size in the Gly73-NLS expressing cells. Data are means ± SEM and analyzed with an unpaired two-tailed Student’s t test (**D**), Pearson’s correlation analysis (**E**), and one-way ANOVA (**B**). ‘‘ns’’ represents non-significant; *p < 0.05, **p < 0.01, and ***p < 0.001.

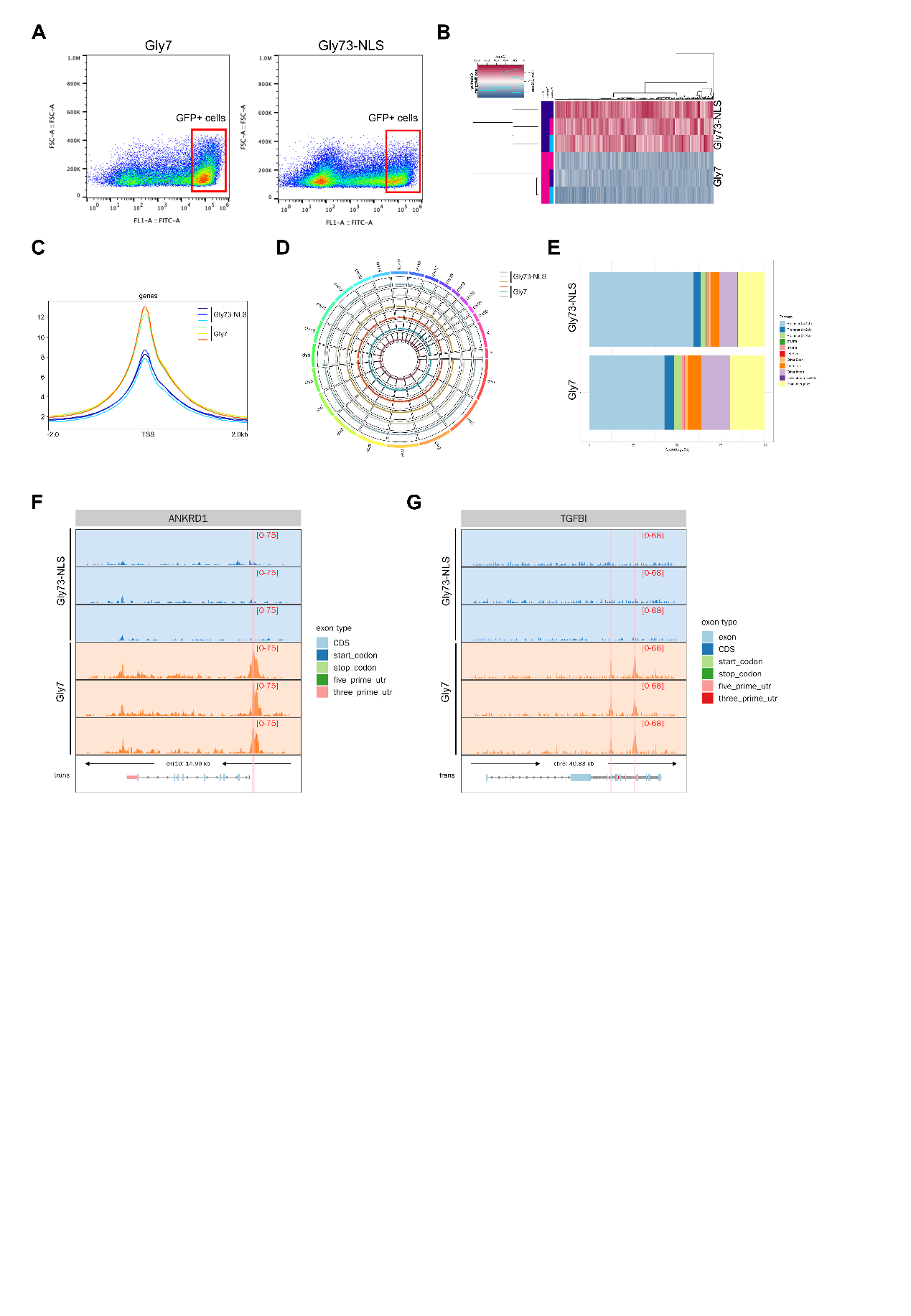
**Supplemental Figure 5. ATAC-seq workflow and additional analyses of chromatin accessibility changes induced by intranuclear polyG aggregation.** **(A)** GFP-high SH-SY5Y cells expressing Gly7 or Gly73-NLS were isolated by flow cytometry and subjected to ATAC-seq analysis. **(B)** Unsupervised clustering of ATAC-seq samples from Gly7 and Gly73-NLS cells. **(C)** TSS-centered accessibility heatmaps showing reduced chromatin accessibility in Gly73-NLS cells relative to Gly7 controls. **(D)** Circular plot showing the chromosomal distribution of ATAC-seq peaks in Gly7 and Gly73-NLS cells. **(E)** Genomic feature annotation of ATAC-seq peaks in Gly7 and Gly73-NLS cells. **(F)** Representative genome browser tracks at the *ANKRD1* locus showing reduced chromatin accessibility in Gly73-NLS cells relative to Gly7 controls.

**(G)** Representative genome browser tracks at the *TGFBI* locus showing reduced chromatin accessibility in Gly73-NLS cells relative to Gly7 controls.

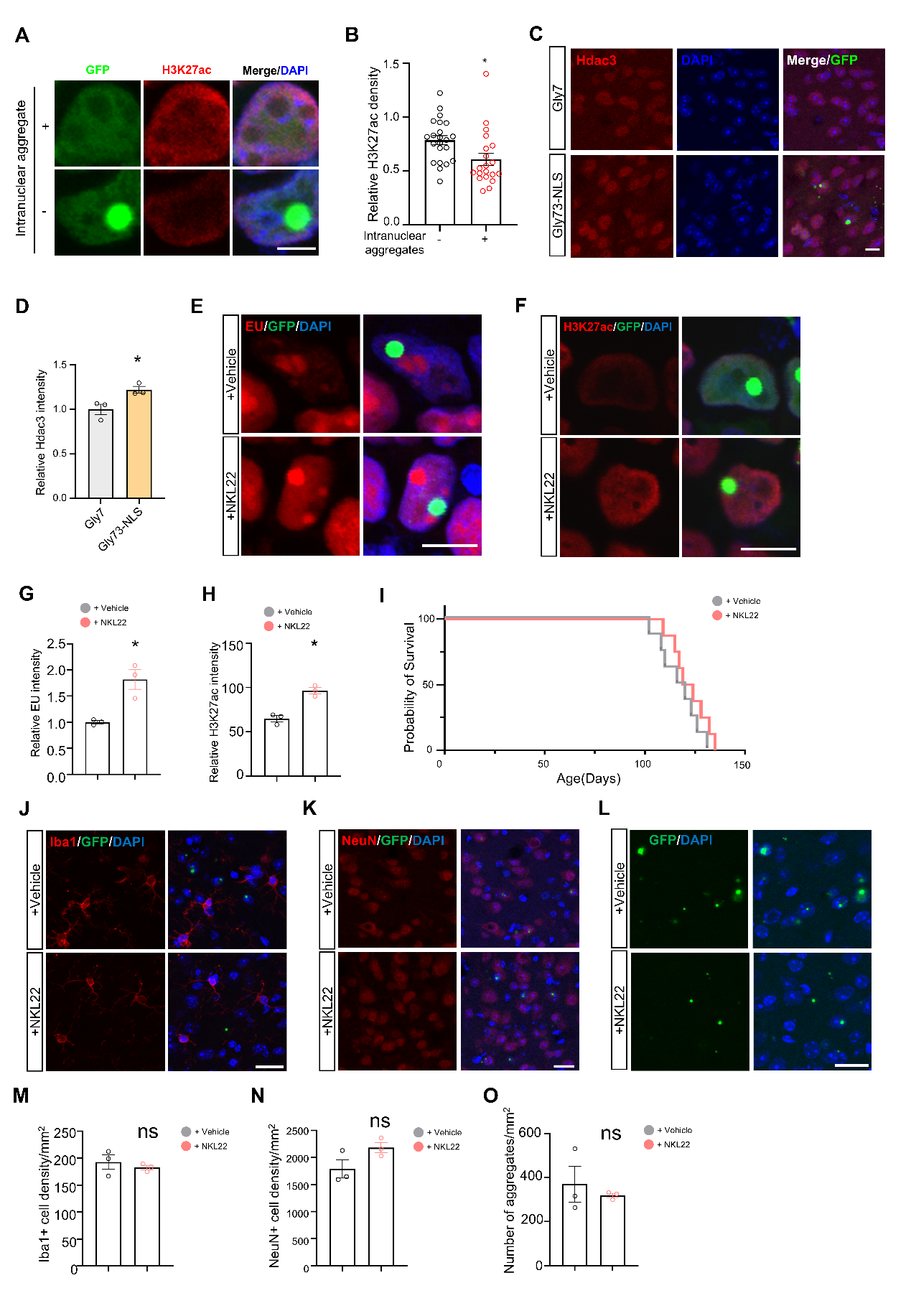
**Supplemental Figure 6. Additional analyses of H3K27ac/HDAC3 changes and the effects of NKL22 in polyG models**. **(A)** Representative images comparing H3K27ac signal in aggregate-negative and aggregate-positive SH-SY5Y cells. Scale bar, 5 μm. **(B)** Quantification of relative H3K27ac intensity in aggregate-negative and aggregate-positive cells (7 cells from each of three independent experiments per group). **(C)** Representative images of HDAC3 immunostaining in mouse brain sections from Gly7 and Gly73-NLS-expressing mice. Scale bar, 5 μm. **(D)** Quantification of relative HDAC3 intensity in the indicated mouse groups (n=3). **(E)** Representative images of EU labeling in polyG-expressing cells treated with vehicle or NKL22. Scale bar, 5 μm. **(F)** Representative images of H3K27ac immunostaining in polyG-expressing cells treated with vehicle or NKL22. Scale bar, 5 μm. **(G)** Quantification of relative EU intensity in vehicle- and NKL22-treated cells (n=3). **(H)** Quantification of relative H3K27ac intensity in vehicle- and NKL22-treated cells (n=3). **(I)** Kaplan–Meier survival curves of Gly73-NLS mice treated with vehicle or NKL22. **(J)** Representative Iba1 immunostaining images in the cerebral cortex of vehicle- and NKL22-treated Gly73-NLS mice. Scale bar, 20 μm. **(K)** Representative NeuN immunostaining images in the cerebral cortex of vehicle- and NKL22-treated Gly73-NLS mice. Scale bar, 20 μm. **(L)** Representative polyG aggregates images in the cerebral cortex of vehicle- and NKL22-treated Gly73-NLS mice. Scale bar, 20 μm. **(M)** Quantification of Iba1-positive cell density (n=3). **(N)** Quantification of NeuN-positive cell density (n=3). **(O)** Quantification of polyG aggregate density (n=3). Data are means ± SEM and analyzed with an unpaired two-tailed Student’s t test (**B, D, G, H, M, N, O**) and log-rank (Mantel–Cox) test (**I**). ‘‘ns’’ represents non-significant; *p < 0.05 and **p < 0.01.

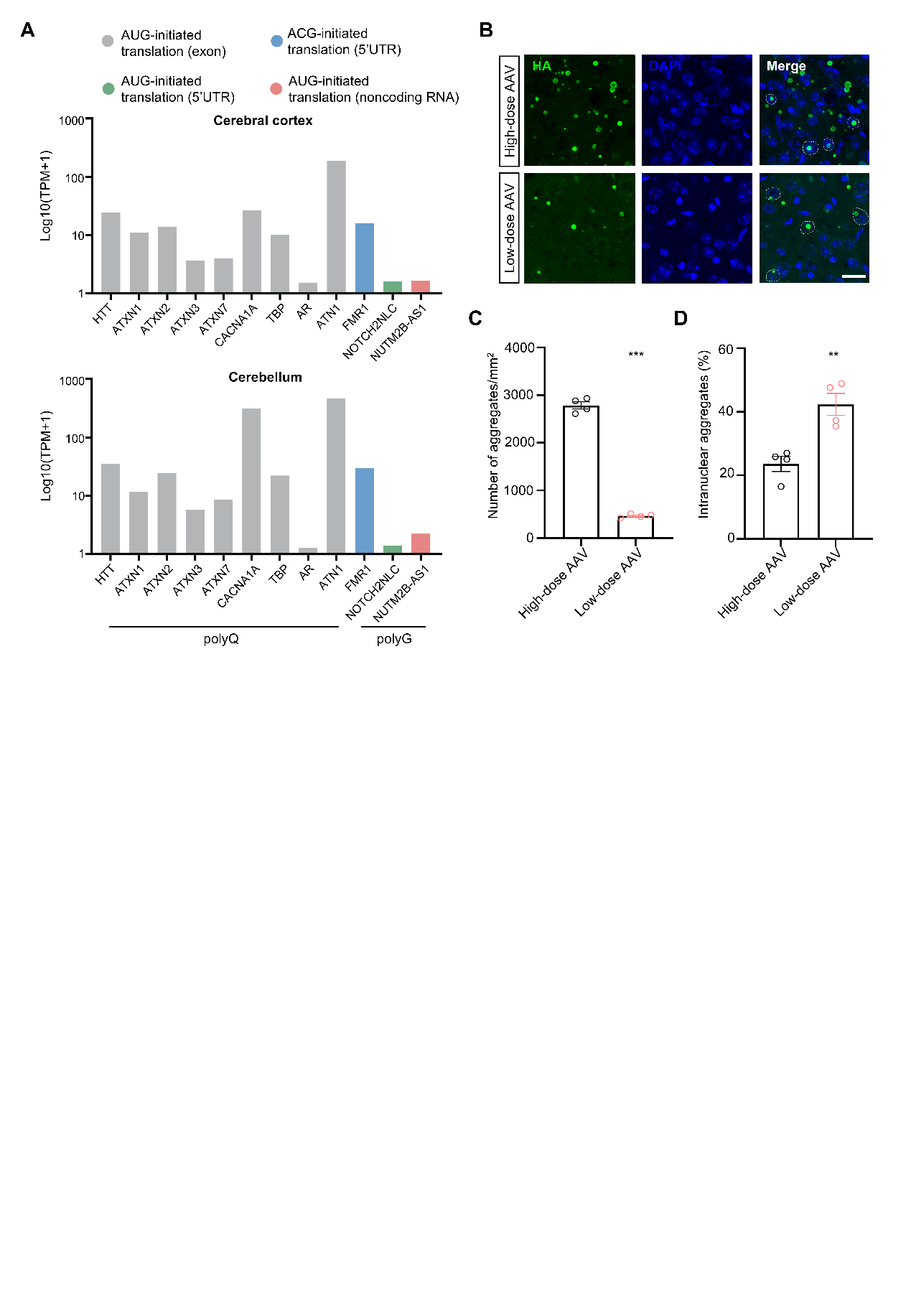
**Supplemental Figure 7. Low expression and small molecular size may contribute to the intranuclear accumulation of polyG proteins**. **(A)** GTEx-based expression levels of genes associated with repeat expansion disorders in the human cerebral cortex and cerebellum. **(B)** Representative images of cortical polyG aggregates in mice injected with low-dose or high-dose AAV-uN2CpolyG-HA. Scale bar, 20 μm. **(C)** Quantification of the number of aggregates in the cerebral cortex of mice injected with low-dose or high-dose AAV (n=4). **(D)** Quantification of the percentage of intranuclear aggregates among total cortical inclusions in the indicated groups (n=4). Data are means ± SEM and analyzed with an unpaired two-tailed Student’s t test (**C, D**). *p < 0.05, **p < 0.01, and ***p < 0.001.

**Table S1. Expression of histone acetylation regulators in the cortex and cerebellum of control and Gly73-NLS mice.**

| Gene | Cortex Gly7 | Cortex  Gly73-NLS | Log2 Fold change | P value | Cerebellum Gly7 | Cerebellum  Gly73-NLS | Log2 Fold change | P value |
| --- | --- | --- | --- | --- | --- | --- | --- | --- |
| *Hdac1* | 8.808 | 8.802 | -0.0008192 | 0.991 | 11.698 | 13.570 | 0.21422049 | 0.032 |
| *Hdac2* | 21.775 | 24.145 | 0.14905174 | 0.271 | 21.722 | 22.530 | 0.05265716 | 0.247 |
| *Hdac3* | 22.462 | 26.565 | **0.24200821** | **0.012** | 19.888 | 23.755 | **0.25636928** | **0.003** |
| *Hdac4* | 4.152 | 3.432 | -0.2747204 | 0.107 | 3.582 | 3.442 | -0.0575101 | 0.459 |
| *Hdac5* | 14.268 | 11.282 | -0.3386458 | 0.007 | 16.008 | 13.265 | -0.2711233 | 0.004 |
| *Hdac6* | 8.675 | 8.02 | -0.1132615 | 0.149 | 9.038 | 9.845 | 0.12346746 | 0.160 |
| *Hdac7* | 5.950 | 6.328 | 0.08874593 | 0.496 | 4.585 | 6.16 | 0.42600862 | 0.003 |
| *Hdac8* | 2.738 | 2.605 | -0.0715756 | 0.446 | 6.228 | 5.235 | -0.2504636 | 0.038 |
| *Hdac9* | 3.285 | 2.650 | -0.309901 | 0.237 | 1.3625 | 1.100 | -0.3087527 | 0.122 |
| *Hdac10* | 3.918 | 3.892 | -0.0092362 | 0.958 | 6.112 | 6.370 | 0.05953081 | 0.724 |
| *Hdac11* | 30.560 | 33.955 | 0.15197949 | 0.058 | 25.805 | 32.430 | 0.32967839 | 0.014 |
| *Sirt1* | 4.650 | 4.862 | 0.06446753 | 0.606 | 10.052 | 10.725 | 0.09342331 | 0.176 |
| *Sirt2* | 19.258 | 24.245 | 0.33226683 | 0.006 | 21.240 | 22.932 | 0.11060987 | 0.099 |
| *Sirt3* | 6.035 | 6.075 | 0.00953064 | 0.844 | 3.9275 | 4.0275 | 0.03627331 | 0.750 |
| *Sirt4* | 9.650 | 8.275 | -0.2217696 | 0.030 | 15.275 | 16.095 | 0.0754402 | 0.466 |
| *Sirt5* | 1.890 | 1.938 | 0.03581008 | 0.708 | 3.332 | 2.995 | -0.1540489 | 0.129 |
| *Sirt6* | 4.005 | 3.772 | -0.0862813 | 0.628 | 4.538 | 5.070 | 0.1600881 | 0.295 |
| *Sirt7* | 8.232 | 7.858 | -0.0672602 | 0.352 | 7.880 | 8.072 | 0.03481991 | 0.706 |
| *Ep300* | 6.030 | 4.718 | -0.3541355 | 0.057 | 9.700 | 9.772 | 0.01074293 | 0.868 |
| *Crebbp* | 6.525 | 5.715 | -0.1912244 | 0.105 | 9.032 | 9.322 | 0.04559154 | 0.441 |
| *Kat2a* | 17.215 | 17.042 | -0.0145292 | 0.876 | 12.530 | 13.605 | 0.11875054 | 0.335 |
| *Kat2b* | 6.408 | 6.142 | -0.0609356 | 0.603 | 30.365 | 31.052 | 0.03230005 | 0.653 |

**Table S2. Genomic locations and estimated molecular weights of pathogenic proteins in repeat expansion-associated neurodegenerative disorders of the central nervous system.**

| **Diseases** | **Gene / locus** | **Disease category** | **Expansion location** | **Pathogenic repeat size** | **Pathogenic protein MW (kDa)** | **Inclusion** |
| --- | --- | --- | --- | --- | --- | --- |
| HD | ***HTT*** | polyQ | Exon1 | >39 | ~355 | N>C |
| SCA1 | ***ATXN1*** | polyQ | Exon8 | >44 | ~92 | N>C |
| SCA2 | ***ATXN2*** | polyQ | Exon1 | >34 | **~143** | C>N |
| SCA3 | ***ATXN3*** | polyQ | Exon | 60–87 | **~42** | **N**>C |
| SCA7 | ***ATXN7*** | polyQ | Exon | >36 | **~100** | **N**>C |
| SCA6 | ***CACNA1A*** | polyQ | Exon47 | 20–33 | **~285** | **C**>N |
| SCA17 | ***TBP*** | polyQ | Exon3 | >49 | ~40 | N |
| SBMA | *A*R | polyQ | Exon1 | >37 | ~105 | N>C |
| **DRPLA** | ***ATN1*** | polyQ | Exon5 | 48–93 | ~130 | N>C |
| FXTAS | ***FMR1*** | polyG | 5’UTR | 55–200 | **~12** | **N** |
| **NIID/OPDM3** | ***NOTCH2NLC*** | polyG | 5’UTR | 60–300 | **~11** | **N** |
| OPML | ***NUTM2B-AS1*** | polyG | Long non-coding RNA | ~700 | **~12** | **N** |

**Abbreviations:** HD, Huntington’s disease; SCA, spinocerebellar ataxia; SBMA, spinal and bulbar muscular atrophy; DRPLA, dentatorubral-pallidoluysian atrophy; FXTAS, fragile X-associated tremor/ataxia syndrome; NIID, neuronal intranuclear inclusion disease; OPDM, oculopharyngodistal myopathy; OPML, oculopharyngeal myopathy with leukoencephalopathy; 5′UTR, 5′ untranslated region; N, nuclear inclusion; C, cytoplasmic inclusion. Pathogenic protein molecular weights were estimated based on repeat lengths of Q60, G100, or A20.
